# Knock-down of hippocampal DISC1 in immune-challenged mice impairs the prefrontal-hippocampal coupling and the cognitive performance throughout development

**DOI:** 10.1101/2020.05.01.070300

**Authors:** Xiaxia Xu, Lingzhen Song, Ileana L. Hanganu-Opatz

**Author notes:** equal contribution (Xiaxia Xu, Lingzhen Song). Corresponding authors: Ileana L. Hanganu-Opatz, Dev. Neurophysiology, Center for Molecular Neurobiology, University Medical Center Hamburg-Eppendorf, Falkenried 94, 20251 Hamburg, Germany, Xiaxia Xu, Dev. Neurophysiology, Center for Molecular Neurobiology, University Medical Center Hamburg-Eppendorf, Falkenried 94, 20251 Hamburg, Germany.

## Abstract

Disrupted-in-Schizophrenia 1 (DISC1) gene represents an intracellular hub of developmental processes and has been related to cognitive dysfunction in psychiatric disorders. Mice with whole-brain DISC1 knock-down show memory and executive deficits as result of impaired prefrontal-hippocampal communication throughout development, especially when combined with early environmental stressors, such as maternal immune activation (MIA). While synaptic dysfunction of layer 2/3 pyramidal neurons in neonatal prefrontal cortex (PFC) has been recently identified as one source of abnormal long-range coupling in these mice, it is still unclear whether the hippocampus (HP) is also compromised during development. Here we aim to fill this knowledge gap by combining *in vivo* electrophysiology and optogenetics with morphological and behavioral assessment of immune-challenged mice with DISC1 knock-down either in the whole brain (GE) or restricted to pyramidal neurons in CA1 area of intermediate/ventral HP (i/vHP) (G_HP_E). Both groups of mice show abnormal network activity, sharp-waves (SPWs) and neuronal firing in CA1 area. Moreover, optogenetic stimulation of CA1 pyramidal neurons fails to activate the local circuits in the neonatal PFC. These deficits that persist until pre-juvenile development are due to dendrite sparsification and loss of spines of CA1 pyramidal neurons. As a long-term consequence, DISC1 knock-down in immune-challenged mice leads to poorer recognition memory at pre-juvenile age. Thus, besides PFC, hippocampal CA1 area has a critical role for the developmental miswiring and long-lasting cognitive impairment related to mental illness.

**Significance Statement:** Developmental miswiring within prefrontal-hippocampal networks has been proposed to account for cognitive impairment in mental disorders. Indeed, during development, long before the emergence of cognitive deficits, the functional coupling within these networks is reduced in mouse models of disease. However, the cellular mechanisms of dysfunction are largely unknown. Here we combine in vivo electrophysiology and optogenetics with behavioral assessment in immune-challenged mice with hippocampus-confined DISC1 knock-down and show that pyramidal neurons in CA1 area are critical for the developmental dysfunction of prefrontal-hippocampal communication and cognitive impairment.

## Introduction

Highly dynamic processes control the wiring of neural circuits during development. At its end, reliable communication between brain areas accounts for the complex behavioral abilities of an adult. For example, the coactivation of prefrontal and hippocampal networks in theta-gamma oscillatory rhythms is critical for the precise information flow in mnemonic and executive tasks (Siapas et al., 2005; Spellman et al., 2015; Backus et al., 2016; Eichenbaum, 2017). This prefrontal-hippocampal coupling emerges early in life, and the unidirectional drive from the CA1 pyramidal neurons boosts the initial entrainment of local circuits across prefrontal layers (Brockmann et al., 2011; Bitzenhofer et al., 2017b; Ahlbeck et al., 2018). The long-range coupling during development has been proposed to critically contribute to the adult function and cognitive abilities. On the flip side, disease-related dysfunction and poor behavioral performance might result from developmental miswiring (Chini et al., 2019).

The maturation of connectivity and functional coupling within the brain is controlled by numerous cell autonomous processes as well as extracellular and environmental factors. Disrupted-in-schizophrenia 1 (DISC1) is an intracellular scaffold protein that has been identified as an intracellular hub of developmental processes, in particular synapse regulation (Narayan et al., 2013). Despite its name, DISC1 is unlikely to be a “genetic” factor causing schizophrenia (Sullivan et al., 2012; Ripke et al., 2013). Instead, DISC1 has been proven to illustrate the relevance of abnormal development for multiple mental conditions, because it orchestrates molecular cascades hypothesized to underlie disease-relevant physiological and behavioral deficits (Cuthbert and Insel, 2013). Mouse models mimicking DISC1 dysfunction have impaired memory and attention as results of disrupted prefrontal-hippocampal circuits (Koike et al., 2006; Clapcote et al., 2007; Kvajo et al., 2011; Crabtree et al., 2017). These deficits are more prominent when environmental stressors, such as MIA after additionally disrupt the Disc1 locus (Cash-Padgett and Jaaro-Peled, 2013; Parks et al., 2017). This might d be due to the fact that mutated DISC1 modulates the basal or poly I:C-induced cytokine production by interfering with glycogen synthase kinase-3 (Beurel et al., 2010).

Since DISC1 controls the developmental molecular cascades, it is likely that its dysfunction decisively contributes to early miswiring. Indeed, recent findings revealed that abnormal DISC1 expression perturbs the maturation of prefrontal-hippocampal coupling (Hartung et al., 2016; Chini et al., 2019). Immune challenged mice with a whole-brain truncated form of DISC1 (i.e. dual-hit mice) have disorganized oscillatory activity as well as weaker coupling and directed interactions between PFC and HP at neonatal age. In contrast, the activity patterns and communication within prefrontal-hippocampal networks as well as the early cognitive abilities were largely unaffected in one-hit genetic (G) (i.e., only DISC1) or environmental (E) (i.e., only MIA) mouse models (Hartung et al., 2016). The early dysfunction of dual-hit mice might result from abnormal activity in either one or both brain areas or from disrupted projections from HP to PFC. Recently, we identified transient synaptic deficits of prefrontal layer 2/3 pyramidal neurons as one mechanism underlying the abnormal prefrontal-hippocampal communication throughout development (Chini et al., 2019; Xu et al., 2019). However, it is unknown whether developmental dysfunction in HP contributes to the disrupted prefrontal-hippocampal communication. Here we address this open question by using *in utero* electroporation (IUE) to selectively knock down DISC1 in CA1 pyramidal neurons of i/vHP during perinatal development in mice exposed to MIA (dual-hit G_HP_E mice). We combined *in vivo* electrophysiology with optogenetics to provide direct evidence for the causal contribution of hippocampal pyramidal neurons to the deficits of prefrontal-hippocampal coupling in G_HP_E mice throughout development.

## Materials And Methods

All experiments were performed in compliance with the German laws and the guidelines of the European Community for the use of animals in research and were approved by the local ethical committee (015/17, 015/18). Timed-pregnant C57BL/6J mice from the animal facility of the University Medical Center Hamburg-Eppendorf were used. The day of vaginal plug detection was defined as gestational day(G) 0.5, whereas the day of birth was defined as postnatal day(P) 0.

### Experimental Design

Multisite extracellular recordings and behavioral testing were performed on pups of both sexes during neonatal development (i.e. P8-P10) as well as during pre-juvenile development (i.e. P16-P23). None of the investigated parameters differed between males and females, thus, data for both sexes were pooled. Three genetically engineered mutant mouse models were investigated. First, heterozygous genetically engineered mutant DISC1 mice carrying a Disc1 allele (Disc1Tm1Kara) on a C57BL6/J background were used. Due to two termination codons and a premature polyadenylation site, the allele produces a truncated transcript (Kvajo et al., 2008). Genotypes were determined using genomic DNA and following primer sequences: forward primer 5’-TAGCCACTCTCATTGTCAGC-3’, reverse primer 5’-CCTCATCCCTTCCACTCAGC-3’. The second used model was Df(16)A^+/-^, carrying a chromosomal engineered 1.3-Mb microdeletion ranging from Dgcr2 to Hira, a segment syntenic to the 1.5-Mb human 22q11.2 microdeletion that encompasses 27 genes. Genotypes were determined using genomic DNA and following primer sequence indicating a knockout: forward primer 5’-ATTCCCCATGGACTAATTATGGACAGG-3’, reverse primer 5’-GGTATCTCCATAAGACAGAATGCTATGC-3’. Third, C57BL/6J mice with DISC1 knock-down confined to HP were engineered through transfection with short-hairpin RNA (shRNA) to DISC1 (5’-GGCAAACACTGTGAAGTGC-3’) under H1 promoter-driven pSuper plasmid. To visualize the transfected neurons, DISC1 shRNA was expressed together with tDimer2 under the control of the CAG promoter (pAAVCAG-tDimer2). Mice were transfected by IUE with DISC1 shRNA at G15.5. All three mouse models were challenged by MIA, using the viral mimetic poly I:C (5mg/kg) injected i.v. into the pregnant dams at gestational day G9.5. The resulting offspring mimicking the dual genetic-environmental etiology of mental disorders were classified in three groups depending on the genetic background: GE mice (whole-brain DISC1 knock-down + MIA), Df16E (Df(16)A^+/-^ + MIA), and G_HP_E mice (hippocampal DISC1 knock-down + MIA). The offspring of wild-type C57BL/6J dams injected at G9.5 with saline (0.9%, i.v.) were transfected with control shRNA and were classified as controls (CON mice).

#### In utero electroporation

Starting one day before and until two days after surgery, timed-pregnant C57BL/6J mice received on a daily basis additional wet food supplemented with 2-4 drops Metacam (0.5 mg/ml, Boehringer-Ingelheim, Germany). At G15.5 pregnant mice were injected subcutaneously with buprenorphine (0.05 mg/kg body weight) 30 min before surgery. The surgery was performed on a heating blanket and toe pinch and breathing were monitored throughout. Under isoflurane anesthesia (induction: 5%, maintenance: 3.5%) the eyes of the dam were covered with eye ointment to prevent damage before the uterine horns were exposed and moistened with warm sterile phosphate buffered saline (PBS, 37°C). Solution containing 1.25 μg/μl DNA [pAAV-CAG-ChR2(E123T/T159C)-2A-tDimer2, or pAAV-CAG-tDimer2, or shRNA to DISC1 together with pAAV-CAG-tDimer2 (molar ratio=3:1)] and 0.1% fast green dye at a volume of 0.75-1.25 μl were injected into the right lateral ventricle of individual embryos using pulled borosilicate glass capillaries with a sharp and long tip. Plasmid DNA was purified with NucleoBond (Macherey-Nagel, Germany). 2A encodes for a ribosomal skip sentence, splitting the fluorescent protein tDimer2 from the opsin during gene translation. To target i/vHP, a tri-polar approach was used (Szczurkowska et al., 2016). Each embryo within the uterus was placed between the electroporation tweezer-type paddles (5 mm diameter, both positive poles, Protech, TX, USA) that were oriented at 90° leftward angle from the midline and a 0° angle downward from anterior to posterior. A third custom build negative pole was positioned on top of the head roughly between the eyes. Electrode pulses (30 V, 50 ms) were applied six times at intervals of 950 ms controlled by an electroporator (CU21EX, BEX, Japan). By these means, neural precursor cells from the subventricular zone, which radially migrate into the i/vHP CA1 area, were transfected. The expression was confined to HP and no neighboring neocortical areas were transfected. Uterine horns were placed back into the abdominal cavity after electroporation. The abdominal cavity was filled with warm sterile PBS (37°C) and abdominal muscles and skin were sutured individually with absorbable and non-absorbable suture thread, respectively. After recovery, pregnant mice were returned to their home cages, which were half placed on a heating blanket for two days after surgery.

#### Surgery for in vivo electrophysiological recordings and light stimulation

For neonatal recordings in non-anesthetized state, 0.5% bupivacain / 1% lidocaine was locally applied on the neck muscles. For pre-juvenile recordings under anesthesia, mice were injected i.p. with urethane (1 mg/g body weight; Sigma-Aldrich, MO, USA) prior to surgery. For both age groups, under isoflurane anesthesia (induction: 5%, maintenance: 2.5%) the head of the pup was fixed into a stereotaxic apparatus using two plastic bars mounted on the nasal and occipital bones with dental cement. The bone above the PFC (0.5 mm anterior to bregma, 0.1-0.5 mm right to the midline), hippocampus (3.5 mm posterior to bregma, 3.5 mm right to the midline) was carefully removed by drilling a hole of <0.5 mm in diameter. After a 10 min recovery period on a heating blanket, mouse was placed into the setup for electrophysiological recording. Throughout the surgery and recording session the mouse was positioned on a heating pad with the temperature kept at 37°C.

#### Electrophysiological recordings

A four-shank electrode (NeuroNexus, MI, USA) containing 4×4 recording sites (0.4-0.8 MΩ impedance, 100 μm spacing, 125 μm intershank spacing) was inserted into the prelimbic subdivision (PL) of PFC. A one-shank optoelectrode (NeuroNexus, MI, USA) containing 1×16 recordings sites (0.4-0.8 MΩ impedance, 50 μm spacing) aligned with an optical fiber (105 mm diameter) ending 200 μm above the top recording site was inserted into CA1 area. A silver wire was inserted into the cerebellum and served as ground and reference electrode. Extracellular signals were band-pass filtered (0.1-9,000 Hz) and digitized (32 kHz) with a multichannel extracellular amplifier (Digital Lynx SX; Neuralynx, Bozeman, MO, USA) and the Cheetah acquisition software (Neuralynx). Spontaneous (i.e. not induced by light stimulation) activity was recorded for 20 min at the beginning of each recording session as baseline activity. The position of recording electrodes in the PL and CA1 area of i/vHP was confirmed post mortem. Wide field fluorescence images were acquired to reconstruct the recording electrode position in brain slices of electrophysiologically investigated pups and to localize tDimer2 expression in pups after IUE. Only pups with correct electrode and transfection position were considered for further analysis. In PL, the most medial shank was inserted to target layer 2/3, whereas the most lateral shank was located into layer 5/6. For the analysis of hippocampal LFP, the recording site located in the pyramidal layer, where SPWs reverse (Bitzenhofer and Hanganu-Opatz, 2014) was selected to minimize any non-stationary effects of large amplitude events. For the analysis of hippocampal firing, two channels above and two channels below the site used for LFP analysis were additionally considered.

#### Light stimulation

Pulsatile (laser on-off, 3 ms-long, 8 Hz) or ramp (linearly increasing power, 3 s-long) light stimulations were performed with an arduino uno (Arduino, Italy) controlled diode laser (473 nm; Omicron, Austria). Laser power was adjusted to trigger neuronal spiking in response to >25% of 3 ms-long light pulses at 8 Hz. Resulting light power was in the range of 20-40 mW/mm^2^ at the fiber tip.

### Behavioral Experiments

The exploratory behavior and recognition memory of CON, G_HP_E and GE mice were tested at pre-juvenile age using previously established experimental protocols (Kruger et al., 2012). Briefly, all behavioral tests were conducted in a custom-made circular white arena, the size of which (D: 34 cm, H: 30 cm) maximized exploratory behavior, while minimizing incidental contact with testing objects (Heyser and Ferris, 2013). The objects used for testing of novelty recognition were six differently shaped, textured and colored, easy to clean items that were provided with magnets to fix them to the bottom of the arena. Object sizes (H: 3 cm, diameter: 1.5-3 cm) were smaller than twice the size of the mouse and did not resemble living stimuli (no eye spots, predator shape). The objects were positioned at 10 cm from the borders and 8 cm from the center of the arena. After every trial the objects and arena were cleaned with 0.1 % acetic acid to remove all odors. A black and white CCD camera (VIDEOR TECHNICAL E. Hartig GmbH, Roedermark, Germany) was mounted 100 cm above the arena and connected to a PC via PCI interface serving as frame grabber for video tracking software (Video Mot2 software, TSE Systems GmbH, Bad Homburg, Germany).

#### Exploratory behavior in the open field (OF)

Pre-juvenile mice (P16) were allowed to freely explore the testing arena for 10 min. Additionally, the floor area of the arena was digitally subdivided in 8 zones (4 center zones and 4 border zones) using the zone monitor mode of the VideoMot 2 analysis software (VideoMot 2, TSE Systems GmbH). The time spent by pups in center and border zones, as well as the running distance and velocity were quantified.

#### Novelty recognition paradigms

All protocols for assessing item recognition memory in P17 mice consisted of familiarization and testing trials (Ennaceur and Delacour, 1988). During the familiarization trial each mouse was placed into the arena containing two identical objects and released against the center of the opposite wall with the back to the objects. After 10 min of free exploration of objects the mouse was returned to a temporary holding cage. Subsequently, the test trial was performed after a delay of 5 min post-familiarization. The mice were allowed to investigate one familiar and one novel object with a different shape and texture for 5 min. Since some mice lost interest to achieve the tasks even before the end of investigation time, object interaction during the first 3 minutes was analyzed and compared between the groups. Discrimination ratio was calculated as (Time spent interacting with novel object – time spent interacting with the familiar object) / (Time spent interacting with novel object + time spent interacting with the familiar object).

In the RR task, tested at P19-20, mice experienced two 10 min-long familiarization trials with two different sets of identical objects that were separated by a delay of 30 min. The second familiarization trial was followed after 5 min by a test trial in which one object used in the first and one object used in the second more recent familiarization trial were placed in the arena at the same positions as during the familiarization trials. Object interaction during the first 3 min was analyzed and compared between the groups. All trials were video-tracked and the analysis was performed using the Video Mot2 analysis software. The object recognition module of the software was used and a 3-point tracking method identified the head, the rear end and the center of gravity of the mouse. Digitally, a circular zone of 1.5 cm was created around each object and every entry of the head point into this area was considered as object interaction. Climbing or sitting on the object, mirrored by the presence of both head and center of gravity points within the circular zone, were not counted as interactions. Discrimination ratios were calculated as (Time spent interacting with more recent object – time spent interacting with less recent object) / (Time spent interacting with more recent object + time spent interacting with less recent object).

### Histology and immunohistochemistry

Histological procedures were performed as previously described (Bitzenhofer et al., 2017b; Oberlander et al., 2019; Xu et al., 2019). Briefly, P8-10 and P20-23 mice were anesthetized with 10% ketamine (aniMedica) / 2% xylazine (WDT) in 0.9% NaCl solution (10 μg/g body weight, i.p.) and transcardially perfused with Histofix (Carl Roth) containing 4% paraformaldehyde. Brains were postfixed in Histofix for 24 h and sectioned coronally at 50 mm (immunohistochemistry) or 100 mm (Sholl and spine analysis). Free-floating slices were permeabilized and blocked with PBS containing 0.8 % Triton X 100 (Sigma-Aldrich, MO, USA), 5% normal bovine serum (Jackson Immuno Research, PA, USA) and 0.05% sodium azide. Subsequently, slices were incubated with mouse monoclonal Alexa Fluor-488 conjugated antibody against NeuN (1:200, MAB377X, Merck Millipore, MA, USA) or the rabbit polyclonal primary antibody against DISC1 (1:250, 40-6800, Thermo Fisher Scientific, MA), followed by 2h incubation with Alexa Fluor-488 goat anti-rabbit IgG secondary antibody (1:500, A11008, Merck Millipore, MA). Slices were transferred to glass slides and covered with Fluoromount (Sigma-Aldrich, MO, USA). Wide field fluorescence images were acquired to reconstruct the recording electrode position and the location of tDimer2 expression. High magnification images were acquired by confocal microscopy (DM IRBE, Leica, Germany) to quantify DISC1 expression. For this, the fluorescence intensity of DISC1 in tDimer2-positive neurons was calculated. All images were similarly processed and analyzed using ImageJ software.

### Neuronal morphological analysis

Microscopic stacks were examined on a confocal microscopy (DM IRBE, Leica Microsystems, Zeiss LSN700 and Olympus FX-100). Stacks were acquired as 2048×2048 pixel images (pixel size, 78 nm; Z-step, 500 nm). Sholl analysis and spine density quantification were carried out in the ImageJ environment. For Sholl analysis, images were binarized (*auto threshold*) and dendrites were traced using the semi-automatical plugin *Simple Neurite Tracer*. The traced dendritic tree was analyzed with the plugin *Sholl Analysis*, after the geometric center was identified using the *blow/lasso* tool. For spine density quantification, the length (*line*) and number of spines (*point picker*) on the dendrite of interest (apical, basal, proximal oblique or secondary apical) were quantified.

### Data Analysis

Data were imported and analyzed offline using custom-written tools in MATLAB software version 7.7 (Mathworks). The data were processed as following: (i) band-pass filtered (500-5000 Hz) to detect MUA as negative deflections exceeding five times the standard deviation of the filtered signals and (ii) low-pass filtered (<1500 Hz) using a third order Butterworth filter before downsampling to 1000 Hz to analyze the LFP. All filtering procedures were performed in a phase-preserving manner. The position of Dil-stained recording electrodes in PL (most medial shank confined to layer 2/3, most temporal shank confined to layer 5/6) and CA1 was confirmed post-mortem by histological evaluation. Additionally, electrophysiological features (i.e. reversal of LFP and high MUA frequency over stratum pyramidale of CA1) were used for confirmation of the exact recording position in HP.

#### Detection of neonatal oscillatory activity

Discontinuous oscillatory events were dected using a previously developed unsupervised algorithm (Cichon et al., 2014) and confirmed by visual inspection. Briefly, deflections of the root-mean-square of band-pass (3-100 Hz) filtered signals exceeding a variance-depending threshold were assigned as network oscillations. The threshold was determined by a Gaussian fit to the values ranging from 0 to the global maximum of the root-mean-square histogram. Only oscillatory events >1 s were considered for further analysis. Time-frequency plots were calculated by transforming the data using the Morlet continuous wavelet.

#### Power spectral density

For power spectral density analysis, 1 s-long windows of network oscillations were concatenated and the power was calculated using Welch’s method with non-overlapping windows. For optical stimulation, we compared the average power during the 1.5 s-long time window preceding the stimulation to the last 1.5 s-long time window of light-evoked activity.

#### Single unit activity

SUA was detected and clustered using klusta (Rossant et al., 2016) and manually curated using phy (https://github.com/cortex-lab/phy). Data were imported and analyzed using custom-written tools in the MATLAB.

#### Firing rate

The firing rate was computed by dividing the total number of spikes by the duration of the analyzed time window.

#### Inter-spike-interval

ISI was calculated at 1 ms resolution in the range of 10-200 ms.

#### Spike-triggered LFP power

Spike-triggered LFP spectra were calculated as

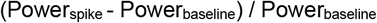

where the spike-triggered power spectrum (Power_spike_) was calculated using Welch’s method for a 200 ms-long time window centered on each spike, and the power spectrum of baseline LFP (Power_baseline_) was averaged for two time windows, 100-300 ms and 200-400 ms before each spike. *Detection of* SPWs *in HP*. The filtered signal (1-300 Hz) was subtracted from the signal recorded 100 μm above and 100 μm below stratum pyramidale. SPWs were detected as peaks above 5 times the standard deviation of the subtracted signal.

#### Phase locking value

PLV was developed to analyze the strength of phase synchronization. Extracting the phase of two signals, PLV was defined as following,

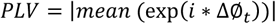

with ΔØ*_t_* stands for phase difference between the two signals at time point t. The value of PLV ranged between 0 (no synchrony) and 1 (max synchrony).

#### Spectral coherence

Coherence was calculated using the coherency method. Briefly, the coherence was calculated (using the functions *cpsd.m* and *pwelch.m*) by cross-spectral density between the two signals and normalized by the power spectral density of each. The computation of the coherence C over frequency (f) for the power spectral density P of signal X and Y was performed according to the formula:

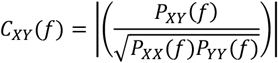

#### Directionality methods

To investigate the directionality of functional connectivity between PFC and HP, gPDC was used. gPDC is based on linear Granger causality measure in the frequency domain. The method attempts to describe the causal relationship between multivariate time series based on the decomposition of multivariate partial coherence computed from multivariate autoregressive models. The LFP signal was divided into 1s-long segments containing the oscillatory activity. After de-noising using MATLAB wavelet toolbox, gPDC was calculated using a previously described algorithm (Baccala and Sameshima, 2001; Baccala et al., 2007).

#### Estimation of light propagation

The spatial pattern of light propagation *in vivo* was estimated using a previously developed model (Stujenske et al., 2015) based on Monte Carlo simulation (probe parameters: light fiber diameter: 50 μm, numerical aperture: 0.22, light parameters: 594 nm, 0.6 mW).

#### Pearson’s correlation

For correlation between gPDC and NOR/RR, we computed Pearson’s correlation using *corrplot.m* in MATLAB.

### Statistical analysis

Statistical analyses were performed in MATLAB environment. Significant differences were detected by paired t-test or one-way ANOVA followed by Bonferroni-corrected post hoc analysis. For Sholl analysis, one-way repeated-measures ANOVA was used. Investigators were blinded to the group allocation when Sholl and spine analyses were performed. Data are presented as mean ± sem. Significance levels of p < 0.05 (*), p < 0.01 (**) or p < 0.001 (***) were tested. Statistical parameters can be found in the main text, tables, and/or in the figure legends.

## Results

### Whole-brain DISC1 knock-down in immune-challenged mice perturbs the patterns of network and spiking activity in neonatal intermediate/ventral HP

Developing prefrontal-hippocampal circuits have been shown to be highly sensitive to the detrimental impact of combined genetic defects and environmental stressors (dual-hit), whereas single-hits usually caused milder phenotypes (Hartung et al., 2016; Oberlander et al., 2019). The mechanisms of abnormal long-range communication and wiring in dual-hit mice are still largely unknown. One possibility is that the maturation of PFC is impaired and therefore, the excitatory drive from the HP does not succeed to entrain the local circuits in beta-gamma frequencies.

Indeed, we recently proved that this is a possible mechanism of developmental dysfunction (Chini et al., 2019; Xu et al., 2019). A second mechanism might be that the hippocampal driving force is decreased as result of abnormal function of developing hippocampal circuits in dual-hit mice.

To test the second hypothesis, we firstly characterized in detail the hippocampal patterns of network and firing activity in immune-challenged mice with whole-brain DISC1 knock-down. For this, we performed extracellular recordings of local field potential (LFP) and multiple-unit activity (MUA) from the CA1 area of i/vHP of awake postnatal day (P) 8-10 CON (n=22) and GE (n=19) mice (*Figure 1A*). As previously reported (Brockmann et al., 2011), discontinuous spindle-shaped oscillations with frequency components peaking in theta band (4-12 Hz) intermixed with irregular low amplitude beta-gamma band components (12-50 Hz) are the dominant pattern of network activity in the CA1 area of both mouse groups (*Figure 1B*). However, their properties significantly differed between CON and GE mice, conferring a highly fragmented appearance of hippocampal oscillations in GE mice (*Figure 1C and D*). Their duration of oscillatory events was significantly shorter in GE mice (CON: 4.46 ± 0.24 s, GE: 3.38 ± 0.18 s, F(1,39)=13.31, *p*=7.7*10^-4, one-way ANOVA) at a comparable occurrence (CON: 8.7 ± 0.30 oscillations/min, GE: 9.47 ± 0.38 oscillations/min, F(1,39)=2.44, *p*=0.126, one-way ANOVA) (*Figure 1C*). Their spectral composition (4-50 Hz) differed between groups, the GE mice having hippocampal events with weaker power when compared to CON (4-50 Hz, F(1,39)=4.29, *p*=0.045, one-way ANOVA) (*Figure 1D*). The overall firing of single neurons in the hippocampal CA1 area of CON and GE was similar both in its rate (log of firing rate, CON: −0.53 ± 0.29; GE: −0.48 ± 0.39; F(1,39)=0.009, *p*=0.924, one-way ANOVA) and temporal organization (i.e. preferred inter-spike interval (ISI) of 125 ms, corresponding to 8 Hz) (*Figure 1E and F*).

**Figure 1.**
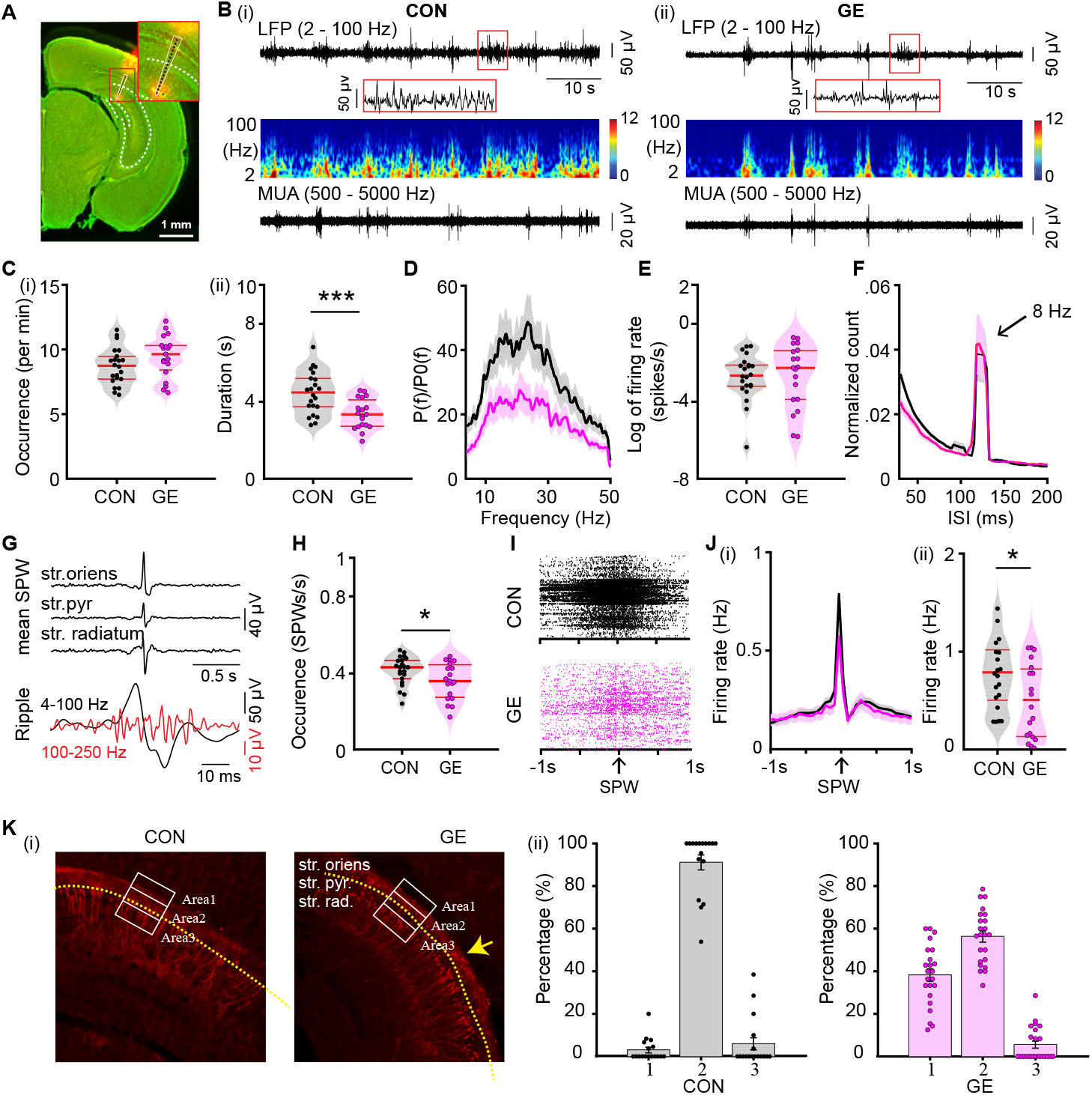
Patterns of network activity and neuronal firing in the CA1 area of i/vHP from neonatal GE mice. **(A)** Digital photomontage reconstructing the location of the DiI-labeled 1×16-site recording electrode (orange) in a 100 μm-thick coronal section containing the CA1 from a P9 mouse. Inset, the position of recording sites (black dots) over the pyramidal layers displayed at higher-magnification. Scale bar, 1mm. **(B)** Extracellular LFP recordings of discontinuous oscillatory activity in the CA1 area from a P9 CON **(i)** and a P9 GE **(ii)** mouse displayed after bandpass (2-100 Hz) filtering (top) and the corresponding MUA after bandpass (500-5000 Hz) filtering (bottom). Traces are accompanied by the color-coded wavelet spectra of the LFP at identical time scale. **(C)** Violin plots displaying the occurrence **(i)** and the duration **(ii)** of hippocampal oscillatory activity recorded in CON and GE mice. **(D)** Averaged power spectra P(f) of discontinuous oscillatory activity normalized to the baseline power P0(f) of time windows lacking oscillatory activity in CON (black) and GE (red) mice. **(E)** Violin plots displaying the firing activity of CA1 neurons in CON and GE mice. **(F)** Histograms of inter-spike intervals for CON (black) and GE (red) mice. Note the prominent inter-spike intervals peak at ~125 ms interval, which corresponds to ~8 Hz. **(G)** Characteristic SPWs and ripple events recorded in the CA1 area. **(H)** Violin plots displaying the occurrence of SPWs in CON and GE mice. **(I)** Examples of spike trains from CA1 neurons aligned to SPWs in CON and GE mice. **(J)** Histograms of spiking activity aligned to SPWs **(i)** and violin plots displaying peak firing rate at SPWs **(ii)** in CON (black) and GE (red) mice. **(K) (i)** Photomicrographs depicting tDimer2-expressing pyramidal neurons (red dots) in the CA1 area of a P9 CON mouse and a P9 GE mouse. The yellow dotted line indicates the pyramidal layer of CA1. The three white blocks with width of 80 μm and length of 200 μm centered on the pyramidal layers correspond to the regions of interest for the quantification of tDimer-transfected neurons. Scale bar, 100 μm. **(ii)** Bar diagram of the distribution of the tDimer-transfected neurons in the three blocks defined in (i) in CON and GE mice. Single data points are represented as dots. Single data points are represented as dots and the red horizontal bars in violin plots correspond to the median and the 25^th^ and 75^th^ percentiles. *p<0.05, ***p<0.001.

Besides spindle-shaped oscillations, prominent SPWs reversing across the pyramidal layer have been recorded in the neonatal CA1 area of CON and GE mice (*Figure 1G*). They were accompanied by ripples (100-250 Hz) and prominent firing. GE mice had fewer SPWs (0.36 ± 0.02 Hz, F(1,39)=4.38, *p*=0.043, one-way ANOVA) when compared with CON mice (0.41 ± 0.02 Hz) (*Figure 1H*). The SPW-related spiking also decreased (CON: 0.76 ± 0.08 Hz, GE: 0.47 ± 0.09, F(1,37)=6.02, *p*=0.019, one-way ANOVA) (*Figure 1I and J*). The rather moderate perturbation of theta oscillations, which have been shown to mainly originate outside CA1 area (Buzsaki, 2002; Janiesch et al., 2011), and the prominent deficits of locally-generated SPWs and related spiking suggest that the i/vHP is compromised in GE mice. Analysis of the position and density of pyramidal neurons suggests that delayed migration of these neurons and correspondingly, perturbed wiring of local circuits, account for the hippocampal dysfunction (*Figure 1 K*).

Since DISC1 is a hub of developmental processes, the hippocampal dysfunction related to a delayed migration of neurons in CA1 area might be specific for the GE mice and not a general feature of mice mimicking the etiology of mental disorders. To test this hypothesis, we performed similar investigations in a dual-hit model that combines MIA with a chromosomal engineered 1.3 - Mb knockout syntenic to the 1.5 - Mb human 22q11.2 microdeletion (Df(16)A^+/-^) (Stark et al., 2008). At P8-10, Df16E mice (n=11) showed similar deficits of hippocampal spindle-burst oscillations (Power, 18.61 ± 4.27 vs. 25.43 ± 7.66, F(1,20)=5.18, *p*=0.035, one-way ANOVA), SPWs (Occurrence, 0.27 ± 0.03 Hz vs. 0.35 ± 0.03 Hz, F(1,20)=5.17, *p*=0.034, one-way ANOVA) and SPW-related firing (0.28 ± 0.08 Hz vs. 0.76 ± 0.16 Hz, F(1,20)=7.69, *p*=0.012, one-way ANOVA) as those reported for GE mice. These data indicate that the hippocampal deficits are a common feature of dual-hit genetic-environmental models of disease.

### DISC1 knock-down in hippocampal pyramidal neurons disturbs the prefrontal oscillatory activity and prefrontal-hippocampal coupling of neonatal immune-challenged mice

When brain-wide expressed, genetic abnormalities, such as DISC1 knock-down or Df(16)A^+/-^ microdeletion, in immune-challenged mice may affect many-fold the communication between limbic areas. To selectively pinpoint their role for hippocampal function, we generated G_HP_E mice in which the DISC1 knock-down was restricted to a lineage of pyramidal neurons in hippocampal CA1 area. For this, we expressed a DISC1 targeting shRNA in the CA1 of the i/vHP by using IUE protocols previously described (Ahlbeck et al., 2018). CON and GE mice received a scrambled/control shRNA instead (*Figure 2A*). The immune challenge of G_HP_E and GE mice was mimicked by MIA with the viral mimetic poly I:C injected at gestational day 9.5. In contrast, CON mice received saline injections at the same age. Staining for NeuN showed that a similar fraction of neurons was transfected in CON (21.89 ± 0.02%; n=6), G_HP_E (20.80 ± 0.01%; n=6) and GE mice (20.59 ± 0.02%; n=6) (*Figure 2B*). The shape of tDimer2-positive neurons and the orientation of primary dendrites confirmed previous findings (Ahlbeck et al., 2018) that the transfection was restricted to cell lineages of pyramidal neurons. In G_HP_E DISC1 was efficiently suppressed by shRNA (*Figure 2C*). The relative DISC1 intensity in CA1 area was significantly weaker (F(1,142)=321.51, *p*=1.06*e-10, one-way ANOVA) in neonatal G_HP_E (40.98 ± 2.56) when compared with CON (101.74 ± 2.14) mice (*Figure 2C*).

**Figure 2.**
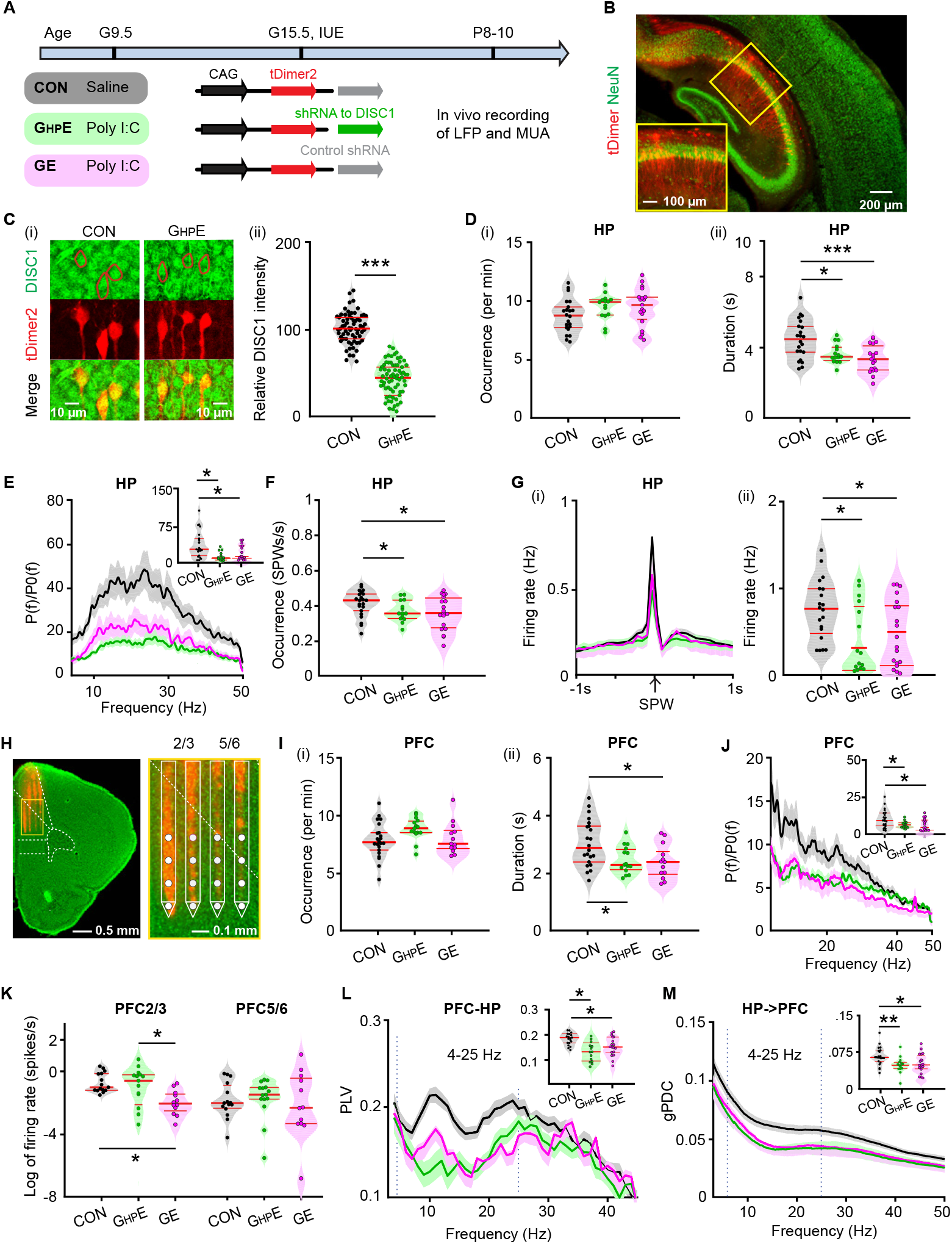
Patterns of network activity and neuronal firing in HP and PFC from neonatal immune challenged mice with HP-confined DISC1 suppression. **(A)** Timeline of experimental protocol and description of the three investigated groups of mice: CON mice, immune-challenged mice with suppression of DISC1 confined to HP (G_HP_E), and immune-challenged mice with brain-wide DISC1 knock-down (GE). For each group the constructs used for IUE to target hippocampal CA1 pyramidal neurons is specified. **(B)** Photomicrographs depicting tDimer2-expressing pyramidal neurons (red) in the CA1 area when stained for NeuN (green) from a P9 mouse. Scale bar, 200 μm. Inset, photograph displaying the tDimer2-expressing cells at a higher magnification. Scale bar, 100 μm. **(C) (i)** Photographs displaying the DISC1 immunoreactivity (green) in relationship with the tDimer2-expression (red) in the CA1 area of i/vHP from P9 G_HP_E and CON mice. Scale bar, 10 μm. **(ii)** Violin plots displaying the relative DISC1 immunoreactivity averaged for G_HP_E and CON mice at P8-P10. **(D)** Violin plots displaying the occurrence **(i)** and the duration **(ii)** of hippocampal oscillatory activity recorded in CON, G_HP_E and GE mice. **(E)** Averaged power spectra P(f) of discontinuous oscillatory activity normalized to the baseline power P0(f) of time windows lacking oscillatory activity in CON (black), G_HP_E (green) and GE (magenta) mice. Insert, violin plots displaying the relative power averaged for 4-50 Hz in CON, G_HP_E and GE mice. **(F)** Violin plots displaying the occurrence of SPWs in the CA1 area of CON, G_HP_E and GE mice. **(G)** Histograms of spiking activity aligned to SPWs **(i)** and violin plots displaying the peak SPW-related firing in CON (black), G_HP_E (green) and GE (magenta) mice **(ii)**. **(H)** Left, digital photomontage reconstructing the location of the DiI-labeled 4 ×4-site recording electrode (orange) in a 100 μm-thick coronal section containing the PFC from a P9 mouse. Scale bar, 0.5 mm. Right, the position of recording sites (white dots) over the prelimbic layers displayed at higher-magnification. Scale bar, 0.1 mm. **(I)** Violin plots displaying the occurrence **(i)** and the duration **(ii)** of prefrontal oscillatory activity recorded in CON, G_HP_E and GE mice. **(J)** Averaged power spectra P(f) of discontinuous oscillatory activity normalized to the baseline power P0(f) of time windows lacking oscillatory activity in CON (black), G_HP_E (green) and GE (magenta) mice. Inset, violin plots displaying the relative power averaged for 4-50 Hz in CON, G_HP_E and GE mice. **(K)** Violin plots displaying the neuronal firing in prefrontal layer 2/3 and layer 5/6 of CON, G_HP_E and GE mice. **(L)** Line plots of mean PLV for oscillatory activity simultaneously recorded in PFC and HP of in CON (black), G_HP_E (green) and GE (magenta) mice. Inset, violin plots displaying the PLV when averaged for 4-25 Hz. **(M)** Line plots of mean gPDC in relationship to frequency for HP→PFC in CON (black), G_HP_E (green) and GE (magenta) mice. Inset, violin plots displaying gPDC when averaged for 4-25 Hz in CON, G_HP_E and GE mice. Single data points are represented as dots and the red horizontal bars in violin plots correspond to the median and the 25^th^ and 75^th^ percentiles. *p<0.05, **p<0.01, ***p<0.001.

To monitor the effects of selective DISC1 knock-down in the i/vHP, we performed extracellular recordings of LFP and MUA from HP of P8-P10 awake CON (n=22), G_HP_E (n=15), and GE mice (n=19). Similar to GE mice, G_HP_E mice showed disorganized oscillatory activity in CA1 area with decreased duration and theta-beta band power (4-50 Hz) but unchanged occurrence of spindle-shaped oscillations (*Table 1, Figure 2D and E*). HP-confined DISC1 knock-down caused reduced SPWs occurrence and SPW-related neuronal firing, similarly to the deficits described for GE mice (*Table 1, Figure 2F and G*).

**Table 1.**
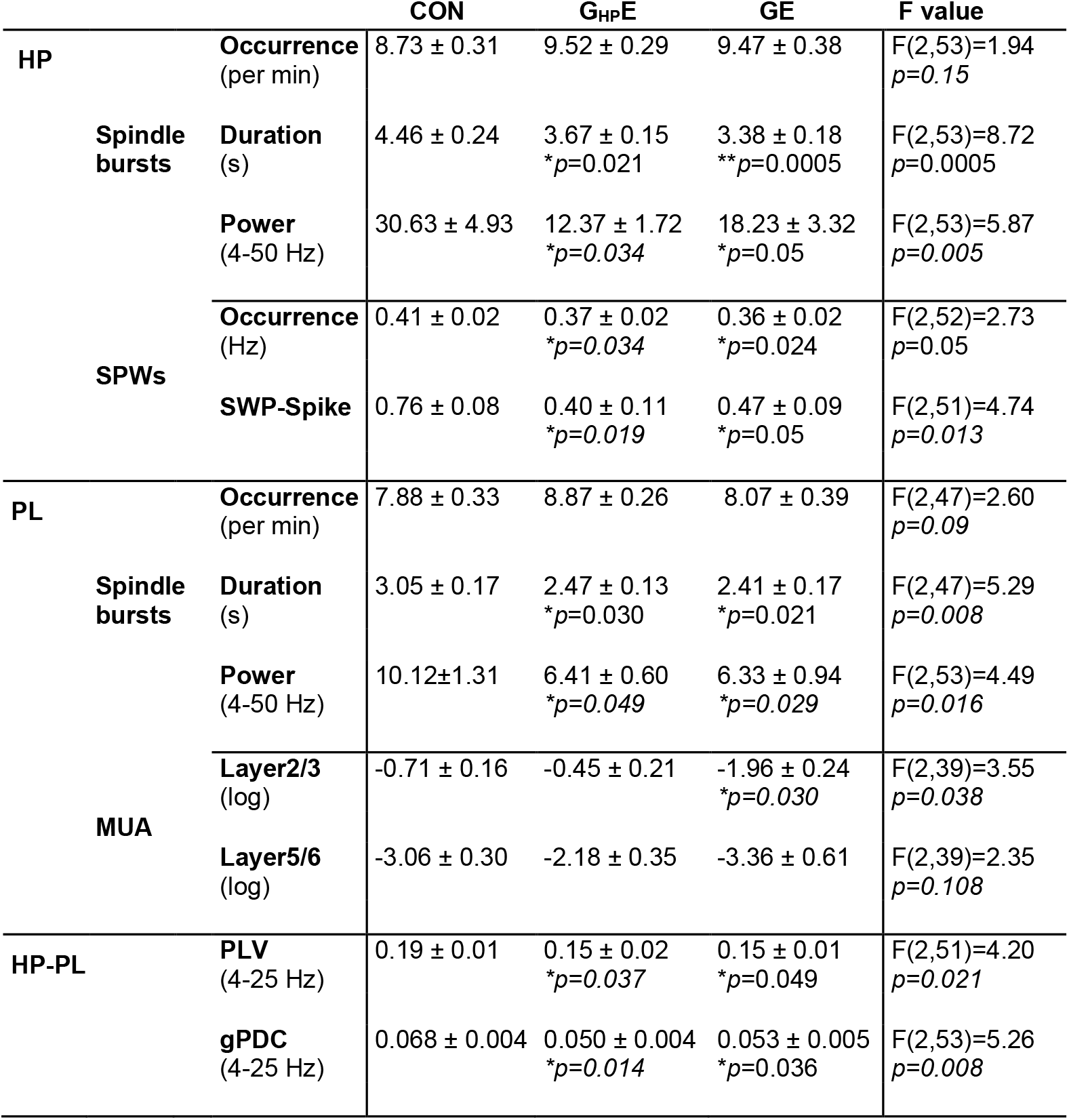
Prefrontal-hippocampal activity patterns and coupling in CON, G_HP_E and GE mice. Data are shown as mean ± SEM. Significance was assessed using one-way ANOVA test followed by Bonferroni-corrected post hoc test and the listed *p* values correspond to comparisons between CON and G_HP_E mice, CON and GE mice.

In light of these findings, the question arises, whether the hippocampal dysfunction in G_HP_E mice is sufficient to affect downstream brain areas with normal DISC1 expression, such as PL. To answer this question, we performed extracellular recordings of LFP and MUA from PL of P8-P10 awake CON (n=22), G_HP_E (n=15) and GE mice (n=19) using four shanks recording electrodes spanning the prelimbic layers 2/3 and 5/6 (*Figure 2H*). In line with previous investigations (Hartung et al., 2016; Chini et al., 2019), the PL of all investigated mice showed discontinuous spindle-shape oscillations with frequencies ranging from theta to beta-low gamma range (20-40 Hz). While the occurrence of these events was similar across the three groups, their duration and power were decreased in GE and G_HP_E mice when compared to CON mice (*Table 1, Figure 2I and J*). As previously reported (Chini et al., 2019), brain-wide DISC1 knock-down in combination with MIA significantly lowered the neuronal firing in prelimbic layer 2/3. In contrast, G_HP_E mice showed normal firing in both layers 2/3 and 5/6 (*Table 1, Figure 2K*) that might result from the sparse hippocampal innervation targeting the PFC. The decreased network entrainment and unchanged firing in PL of G_HP_E mice suggest that the local prelimbic circuits are indirectly impaired, most likely through a weaker drive from HP, which at this age is the main source of PL activation (Brockmann et al., 2011; Ahlbeck et al., 2018). To test this hypothesis, we firstly assessed the synchrony between PL and HP in CON (n=22), G_HP_E (n=15) and GE mice (n=19) by calculating the phase locking values (PLV) that, relying on oscillatory phase information, are not biased by different amplitudes of activity in PFC and HP. In line with previous data (Hartung et al., 2016; Xu et al., 2019), a tight theta-beta band (4-25 Hz) coupling of spindle-bursts between PL and HP has been detected in neonatal CON mice. In contrast, the PLV was significantly lower in G_HP_E and GE mice (F(2, 51)=4.20, *p*=0.021, one-way ANOVA) (*Table 1, Figure 2L*). The directionality of interactions between PL and HP was also affected by both brain-wide and HP-confined DISC1 knock-down in immune-challenged mice. Calculation of generalized partial directed coherence (gPDC), a measure that reflects the directionality of network interactions in different frequency bands, confirmed the prominent drive from HP to PL. In both GE and G_HP_E this drive decreased within 4-25 Hz frequencies (*Table 1, Figure 2M*). These results give first evidence that the suppression of DISC1 restricted to HP has detrimental effects on the function of downstream PL.

### Hippocampal dysfunction through DISC1 suppression is sufficient to reduce the oscillatory entrainment of PL and prelimbic-hippocampal coupling in neonatal immune-challenged mice

To add causal evidence to the hypothesis that the hippocampal dysfunction is critical for the abnormal coupling between PL and HP, we selectively transfected the hippocampal pyramidal neurons in CON, G_HP_E and GE mice with a highly efficient fast-kinetics double mutant ChR2E123T/T159C (ET/TC) (Berndt et al., 2011) and the red fluorescent protein tDimer2 by IUE (*Figure 3A*). For G_HP_E mice, constructs coding for ChR2E123T/T159C (ET/TC) were transfected together with shRNA to DISC1. For selective targeting i/vHP, a previously developed protocol for IUE using three paddles was used (Szczurkowska et al., 2016; Ahlbeck et al., 2018). This method enables stable area and cell type-specific transfection of hippocampal neurons already prenatally without the need of cell-type specific promotors of a sufficiently small size (Baumgart and Grebe, 2015; Szczurkowska et al., 2016) (*Figure 3Bi*). The expression rate and distribution of tDimer2-positive neurons within the iso-contour lines for light power of 1 mW/mm^2^ were similar in CON (0.134 ± 0.009 / 1000 μm^2^, n=16), G_HP_E (0.127 ± 0.008 / 1000 μm^2^, n=15) and GE (0.126 ± 0.010 / 1000 μm^2^, n=18) (F(2, 46)=0.224, *p*=0.800, one-way ANOVA) (*Figure 3Bii*). As previously shown, the transfection procedure had no effects on the overall development of mice (weight, somatic development, reflexes) (Ahlbeck et al., 2018).

**Figure 3.**
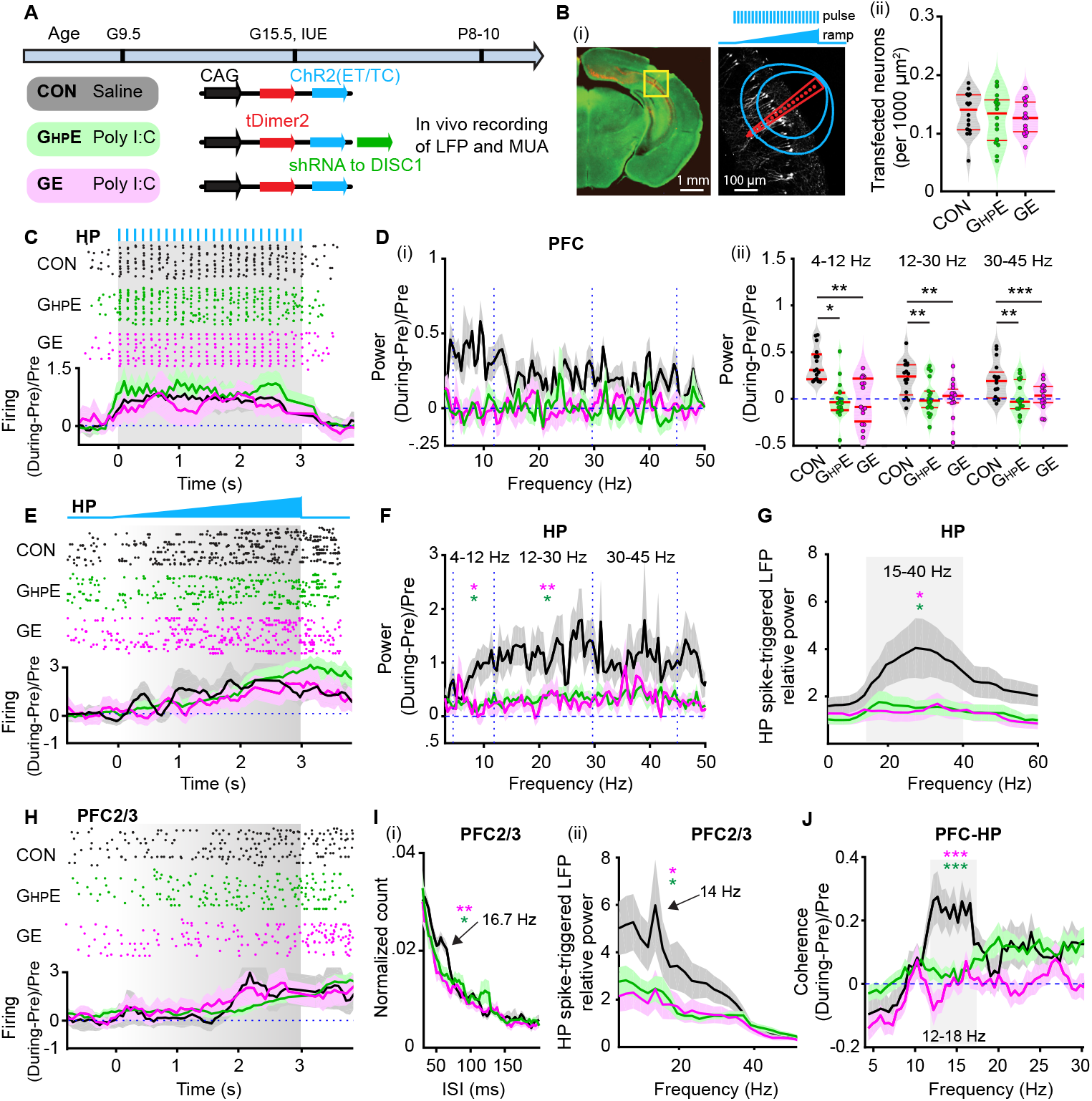
Light-induced activation of i/vHP and prefrontal-hippocampal coupling in immune challenged mice with whole-brain and HP-confined DISC1 suppression. **(A)** Timeline of experimental protocol and description of the three investigated groups of mice: CON, G_HP_E and GE. For each group, the constructs used for IUE to target hippocampal CA1 pyramidal neurons is specified. **(B) (i)** Left, ChR2(ET/TC)-tDimer2-expressing cells (red) in a 50 μm-thick Nissl-stained (green) coronal section including CA1 area from a P9 mouse. Scale bar, 1 mm. Right, recording sites together with transfected neurons (white dots). Green lines correspond to the iso-contour lines for light power of 1 and 10 mW/mm^2^, respectively. 3 s-long pulse (8 Hz) and ramp stimulation are applied to activate hippocampal pyramidal neurons. Scale bar, 100 μm. **(ii)** Violin plots displaying the number of transfected neurons within the iso-contour lines for light power of 1 mW/mm^2^. **(C)** Top, representative raster plot of hippocampal SUA in response to 8 Hz pulse stimulation (3 ms-long pulse, 473 nm) in the CA1 area of P9 CON, G_HP_E and GE mice. Bottom, Histograms of hippocampal firing activity during 8 Hz pulse stimulation normalized to the activity before stimulation in CON (black), G_HP_E (blue) and GE (red) mice. **(D) (i)** Power of prefrontal oscillatory activity during pulse stimulation of CA1 pyramidal neurons normalized to the activity before stimulation in CON (black), G_HP_E (green) and GE (magenta) mice. **(ii)** Violin plots displaying the oscillatory power averaged for different frequency bands (4–12 Hz, 12–30 Hz, 30– 45 Hz) in response to pulse stimulation for all investigated mice. **(E)** Top, representative raster plot of hippocampal spiking in response to ramp stimulation (3 s duration, 473 nm) in CON, G_HP_E and GE mice. Bottom, Histograms of hippocampal firing activity during ramp stimulation normalized to the activity before stimulation in CON (black), G_HP_E (green) and GE (magenta) mice. **(F)** Power of hippocampal oscillatory activity during ramp stimulation of CA1 pyramidal neurons normalized to the activity before stimulation in CON (black), G_HP_E (green) and GE (magenta) mice. **(G)** Line plots of frequency-dependent relative power of spike-triggered LFP in HP of CON (black), G_HP_E (green) and GE (magenta) mice. **(H)** Top, representative raster plot of prefrontal spiking in response to ramp stimulation (3s duration, 473 nm) in HP in one CON, G_HP_E and GE mouse. Bottom, histograms of prefrontal firing activity during ramp stimulation normalized to the activity before stimulation in CON (black), G_HP_E (green) and GE (magenta) mice. **(I) (i)** Histograms of inter-spike intervals for layer 2/3 prefrontal neurons during ramp stimulation of hippocampal CA1 pyramidal neurons for CON (black), G_HP_E (green) and GE (magenta) mice. Note the prominent inter-spike intervals peak at ~16.7 Hz. **(ii)** Line plots of frequency-dependent relative power of hippocampal spike-triggered LFP in prefrontal layer 2/3 of CON (black), G_HP_E (green) and GE (magenta) mice. **(J)** Line plots of coherence between PFC and HP during ramp stimulation of hippocampal CA1 pyramidal neurons normalized to coherence values before stimulation. Single data points are represented as dots and the red horizontal bars in violin plots correspond to the median and the 25^th^ and 75^th^ percentiles. *p<0.05, **p<0.01, ***p<0.001. Magenta stars correspond to the comparison between GE and CON mice. Green stars correspond to the comparison between G_HP_E and CON mice.

First, we assessed the efficiency of light stimulation in evoking action potentials in hippocampal pyramidal neurons *in vivo*. For this, we stimulated the i/vHP with pulsed blue light (473 nm, 20-40 mW/mm^2^) at 8 Hz, since this frequency has been shown to optimally drive the PL (Ahlbeck et al., 2018). The used light power did not cause local tissue heating that might interfere with neuronal spiking (Stujenske et al., 2015; Bitzenhofer et al., 2017a). In all three mouse groups, light stimulation induced precisely timed firing (latency < 10 ms) of CA1 neurons (*Figure 3C*).

Second, to investigate the effects of hippocampal activation on downstream prelimbic circuits, we performed extracellular recordings of LFP in the PL during pulsed light stimulation of CA1 area in CON (n=15), G_HP_E (n=18) and GE (n=15) mice (*Figure 3D*). In CON mice, the light-induced hippocampal firing significantly augmented the prefrontal oscillatory activity in all frequency bands, as reflected by the higher power during stimulation when compared with the time window before the train of pulses (*Table 2, Figure 3D*). In contrast, the light-induced hippocampal firing failed to boost the prelimbic oscillatory activity in G_HP_E and GE mice (*Figure 3D*).

**Table 2.** Prefrontal-hippocampal activity patterns and coupling induced by pulsed or ramp light stimulation of ChR2(ET/TC)-transfected pyramidal neurons in the CA1 area of i/vHP. Data are shown as mean ± SEM. Significance was assessed using one-way ANOVA test followed by Bonferroni-corrected post hoc test and the listed *p* values correspond to comparisons between CON and G_HP_E mice, CON and GE mice.

|  |  |  | CON | G<sub>HP</sub>E | GE | F values |
| --- | --- | --- | --- | --- | --- | --- |
| Activating HP by pulses stimulation |  |  |  |  |  |  |
| PFC | Power<br>(During-pre)/pre | 4-12 Hz | 0.234±0.049 | 0.059 ± 0.042<br>*\*p=0.016* | 0.023 ± 0.047<br>*\*\*p=0.005* | F(2,45)=6.47<br>*p=0.0034* |
|  |  | 12-30 Hz | 0.210±0.049 | 0.045 ± 0.032<br>*\*\*p=0.0096* | -0.007 ± 0.039<br>*\*\*p=0.0011* | F(2,45)=8.21<br>*p=0.0009* |
|  |  | 30-45 Hz | 0.184±0.035 | 0.051 ± 0.027<br>*\*\*p=0.007* | 0.004 ± 0.030<br>*\*\*\*p=0.0004* | F(2,45)=9.38<br>*p=0.0004* |
| Activating HP by ramp stimulation |  |  |  |  |  |  |
| HP | Power<br>(During-pre) / pre | 4-12 Hz | 0.186±0.045 | 0.070 ± 0.031<br>*\*p=0.011* | 0.026 ± 0.050<br>*\*p=0.024* | F(2,45)=3.98<br>*p=0.026* |
|  |  | 12-30 Hz | 0.267±0.051 | 0.091 ± 0.043<br>*\*p=0.018* | 0.046 ± 0.043<br>*\*\*p=0.004* | F(2,45)=6.62<br>*p=0.003* |
|  |  | 30-45 Hz | 0.224±0.054 | 0.102 ± 0.046 | 0.089 ± 0.049 | F(2,45)=2.34<br>*p=0.108* |
|  | Spike-triggered LFP | 15-40 Hz | 3.08±0.59 | 1.43 ± 0.27<br>*\*p=0.018* | 1.42 ± 0.40<br>*\*p=0.021* | F(2,43)=5.26<br>*p=0.009* |
| PFC2/3 | Firing interval |  | 0.020±0.001 | 0.015 ± 0.002<br>*\*p=0.04* | 0.014 ± 0.002<br>*\*\*p=0.005* | F(2,39)=6.37<br>*p=0.004* |
|  | HP spike-triggered LFP (15-40 Hz) |  | 7.06±2.19 | 3.03 ± 0.72<br>*\*p=0.044* | 2.90 ± 0.88<br>*\*p=0.042* | F(2,45)=3.22<br>*p=0.049* |
| PFC-HP | Coherence | 12-18 Hz | 0.220±0.044 | 0.041 ± 0.023<br>*\*\*\*p=0.0003* | -0.004 ± 0.024<br>*\*\*\*p=0.00001* | F(2,41)=15.33<br>*p=0.000001* |

The weaker hippocampal drive to PL in G_HP_E mice might result from abnormal network entrainment of the i/vHP. To test this hypothesis, we applied ramp stimulations that, in contrast to light pulses, trigger more physiological firing and do not induce power contamination by repetitive and large voltage deflections (Bitzenhofer et al., 2017a). Ramp stimulation (3 s duration) of CA1 neurons led to sustained increase of spike discharge and augmented theta-beta oscillatory power in the HP of CON mice (*Table 2, Figure 3E*). These effects were absent in GE and G_HP_E mice. Moreover, despite similar hippocampal firing responsiveness to light stimuli, the ability of CA1 neurons contributing to network oscillations in beta-gamma frequencies (20-40 Hz) dramatically decreased in GE and G_HP_E mice as shown by the weaker spike-triggered LFP relative power (*Table 2, Figure 3G*).

Consistent with the excitatory drive from HP to PL during neonatal development, ramp stimulation-induced CA1 firing was relayed to PL and caused augmentation of prelimbic firing across layers in all investigated mouse groups (*Figure 3H*). Since the axonal projections of CA1 neurons target prelimbic layer 5/6 neurons, the firing increase in these layers was stronger than in the layer 2/3 (2.97 ± 0.31 vs. 1.52 ± 0.25, F(1, 16)=14.75, *p*=0.0014, one-way ANOVA), yet lacked temporal coordination in all mice. In contrast, the firing within prelimbic layer 2/3 in CON induced by ramp stimulation of CA1 pyramidal neurons had a preferred inter-spike interval of ~60 ms, equivalent to a population firing at 16.7 Hz (*Figure 3Ii*). Correspondingly, the HP spike-triggered prelimbic LFP relative power peaked at similar frequencies (*Figure 3Iii*). These peaks were absent in GE and G_HP_E mice, reflecting abnormal entrainment of prelimbic circuits. Moreover, ramp-induced activation of CA1 pyramidal neurons boosted the synchrony between PL and HP in CON mice in a frequency-specific manner (peak at 12-18 Hz), yet it did not induce a coherence increase in mice with whole-brain DISC1 suppression (*Table 1, Figure 3J*). Even when the DISC1 suppression is confined to HP, the coherence did not significantly increase during ramp stimulus and augmented mainly after it, most likely as result of non-specific network boosting.

Taken together, these results indicate that hippocampal DISC1 suppression in combination with MIA lead to CA1 dysfunction that, on its turn, causes abnormal coupling within neonatal prefrontal-hippocampal networks.

### DISC1 knock-down causes major morphological and synaptic deficits of pyramidal neurons in CA1 area

The abnormal firing and oscillatory entrainment of HP and consequently, the weaker prelimbic-hippocampal coupling in GE and G_HP_E mice might relate to abnormal morphology and connectivity of CA1 pyramidal neurons. To test this hypothesis, we undertook a detailed histological examination of the cytoarchitecture of tDimer-labeled hippocampal pyramidal neurons of P10 CON, G_HP_E and GE mice (n=17~19 neurons from 3 mice in each group). The complexity of dendritic branching was assessed by Sholl analysis of three-dimensionally reconstructed hippocampal pyramidal neurons. When compared with CON mice, both G_HP_E and GE mice showed a highly significant reduction of dendritic branching of hippocampal pyramidal neurons (condition effect, p<1*10^-9^) (*Figure 4A*). These deficits were particularly prominent within a radius of 20-150 μm from the cell soma center. Next, we examined the spine density along the dendrites of hippocampal pyramidal neurons. In G_HP_E and GE mice the density of dendritic spines was significantly lower when compared with CON mice (*Figure 4 B, C*). The magnitude of reduction was similar for basal dendrites (F(2, 49)=10.21, *p*=1.96*10^-4^, one-way ANOVA), proximal oblique dendrites (F(2, 50)=9.31, *p*=3.66*10^-4^, one-way ANOVA), and secondary apical dendrites (F(2, 48)=8.14, *p*=9.03*10^-4^, one-way ANOVA). Thus, CA1 pyramidal neurons in G_HP_E and GE mice show simplified dendritic arborization and decreased spine density. As a consequence, the hippocampal activity, especially the excitability of individual neurons and the locally-generated SPWs, might be perturbed and the coupling with downstream prelimbic neurons, diminished.

**Figure 4.**
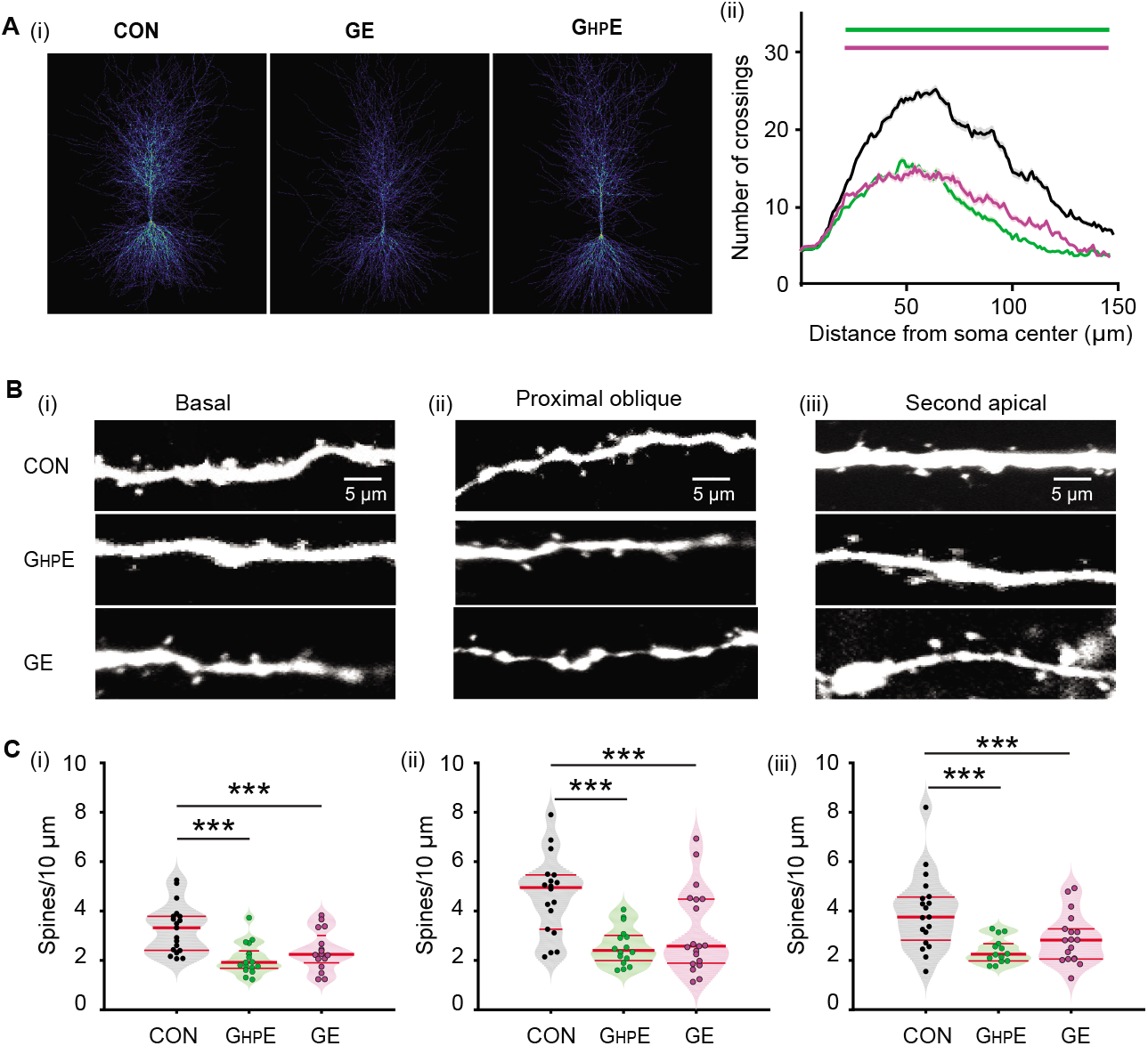
Morphology of hippocampal pyramidal neurons in neonatal G_HP_E and GE mice. **(A) (i)** Heatmap displaying an overlay of all traced dendrites of transfected CA1 pyramidal neurons in CON, G_HP_E and GE mice. **(ii)** Graph displaying the average number of dendritic intersections within a 150 μm radius from the soma center of CA1 pyramidal neurons in CON (black, n=21 neurons from 3 mice), G_HP_E (blue, n=21 neurons from 3 mice) and GE (red, n=21 neurons from 3 mice) mice. Green and magenta bars indicate significant difference (***p<0.001) between CON and G_HP_E mice and between CON and GE mice, respectively. **(B)** Photograph displaying representative basal **(i)**, proximal oblique **(ii)** and second apical dendrites **(iii)** of CA1 pyramidal neurons from a P9 CON, a P9 G_HP_E and a P9 GE mouse. Scale bar, 5 μm. **(C)** Violin plots displaying the spine density on basal **(i)**, proximal oblique **(ii)**, and second apical dendrites **(iii)** of CA1 pyramidal neurons from CON (20 neurons from 3 mice), G_HP_E (20 neurons from 3 mice) and GE (21 neurons from 3 mice) mice. Single data points are represented as dots and the red horizontal bars in violin plots correspond to the median and the 25^th^ and 75^th^ percentiles. ***p<0.001.

### Hippocampal DISC1 knock-down in immune-challenged mice causes neuronal and network deficits as well as cognitive impairment at pre-juvenile age

To test the long-term consequences of hippocampal dysfunction and abnormal prefrontal-hippocampal coupling, we investigated CON, GE, and G_HP_E mice at pre-juvenile age (P20-P23) (*Figure 5A*). The DISC1 suppression persisted, yet at a lower magnitude, until this age, as revealed by the significantly (F(1,146)=25.323, *p*=4.85*10^-7^, one-way ANOVA) lower DISC1 expression in G_HP_E (n=73 neurons) when compared with CON (n=75 neurons) mice (*Figure 5B*). Of note, CA1 pyramidal neurons seem to respond to IUE shRNA suppression differently compared with prelimbic neurons, where the DISC1 knock-down was temporally restricted to the neonatal period (Xu et al., 2019).

**Figure 5.**
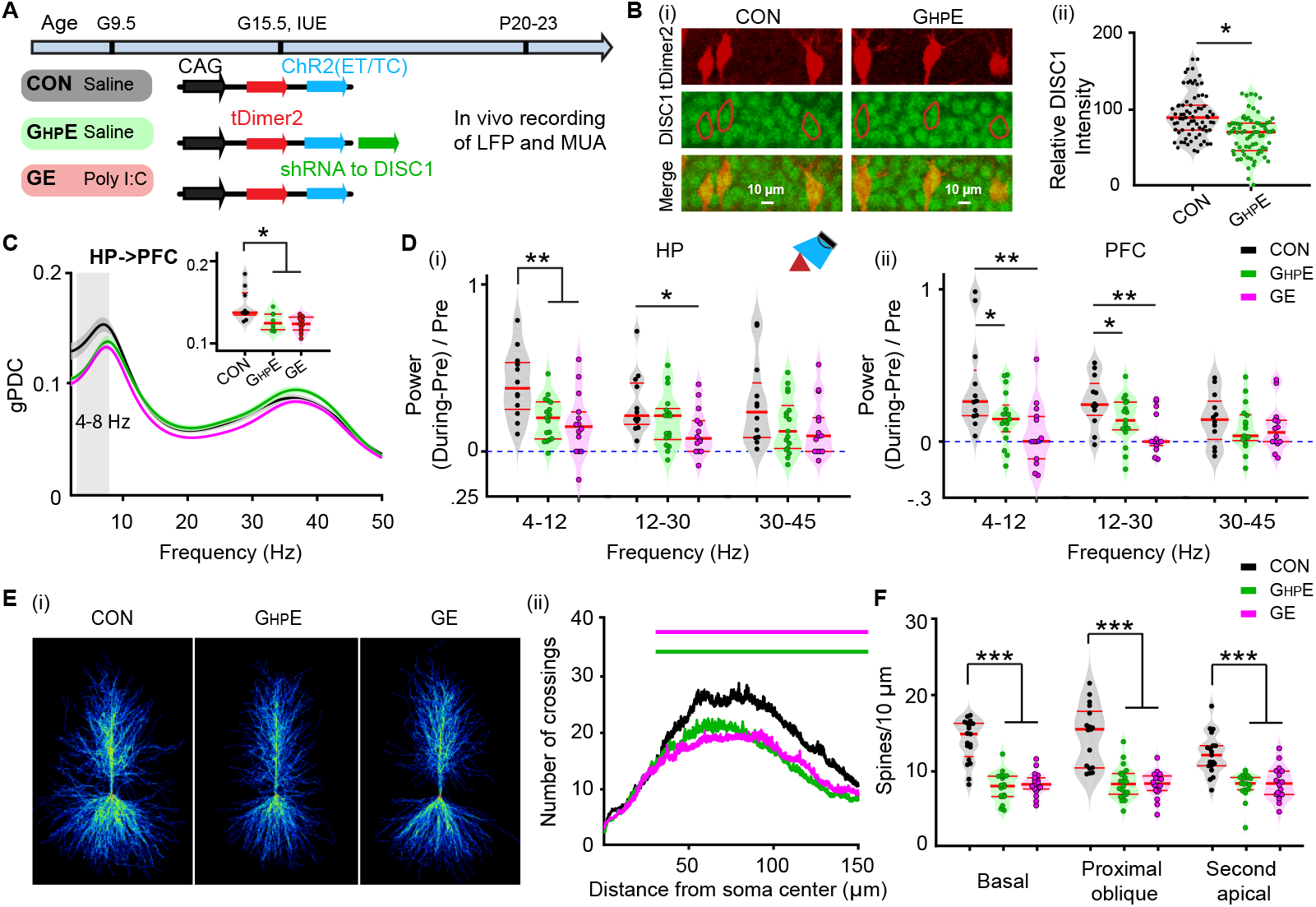
Patterns of pre-juvenile network and neuronal firing in the CA1 area of i/vHP of immune challenged mice with whole-brain or HP-confined DISC1 suppression. **(A)** Timeline of experimental protocol and description of the three investigated groups of mice: CON, G_HP_E and GE. For each group the constructs used for IUE to target hippocampal CA1 pyramidal neurons is specified. **(B) (i)** Photographs displaying the DISC1 immunoreactivity (green) in relationship with the tDimer2-expression (red) in the CA1 area of i/vHP of P21 G_HP_E and CON mice. Scale bar, 10 μm. **(ii)** Violin plots displaying the relative DISC1 immunoreactivity averaged for G_HP_E and CON mice at P20-P23. **(C)** Line plots of mean gPDC in relationship to frequency for HP→PFC in CON (black), G_HP_E (green) and GE (magenta) mice. Inset, violin plots displaying gPDC when averaged for 4-8 Hz in CON, G_HP_E and GE mice. **(D)** Violin plots displaying the hippocampal **(i)** and prefrontal **(ii)** oscillatory power averaged for different frequency bands (4–12 Hz, 12–30 Hz, 30–45 Hz) in response to ramp stimulation in HP for all investigated mice. **(E) (i)** Heatmap displaying an overlay of all traced dendrites of transfected CA1 pyramidal neurons in CON, G_HP_E and GE mice. **(ii)** Graph displaying the average number of dendritic intersections within a 150 μm radius from the soma center of CA1 pyramidal neurons in CON (black, n=21 neurons from 3 mice), G_HP_E (blue, n=21 neurons from 3 mice) and GE (red, n=21 neurons from 3 mice) mice. Green and magenta bars indicate significant difference (***p<0.001) between CON and G_HP_E mice and between CON and GE mice, respectively. **(F)** Violin plots displaying the spine density on basal, proximal oblique, and second apical dendrites of CA1 pyramidal neurons from CON, G_HP_E and GE mice. Single data points are represented as dots and the red horizontal bars in violin plots correspond to the median and the 25^th^ and 75^th^ percentiles. *p<0.05, **p<0.01, ***p<0.001.

First, we performed multi-site extracellular recordings of LFP from PL and hippocampal CA1 area of urethane-anesthetized P20-23 CON (n=12), G_HP_E (n=9), and GE (n=15) mice. All investigated mice showed continuous large-amplitude slow rhythms that were superimposed with oscillatory activity in theta (4-12 Hz) and gamma (30-100 Hz) frequencies. These patterns of network activity correspond to the sleep-like rhythms mimicked by urethane anesthesia (Wolansky et al., 2006; Clement et al., 2008). While the impairment of neuronal firing, sharp-waves, and network oscillations was less pronounced at pre-juvenile age when compared with the deficits at neonatal age, the directed interactions between PFC and HP were still compromised (*Figure 5C*). Both GE (0.123 ± 0.003, *p*=0.016, ANOVA followed by Bonferroni-corrected post-hoc test) and G_HP_E (0.127 ± 0.004, *p*=0.002, ANOVA followed by Bonferroni-corrected post-hoc test) mice had smaller gPDC peaks within theta band (4-8 Hz) and thus, weaker drive from HP to PFC, when compared to CON mice (0.147 ± 0.007) mice. Light stimulation of ChR2(ET/TC)-expressing CA1 neurons augmented the power of network oscillations in theta-beta range for CON (theta: 0.404 ± 0.096; beta: 0.283 ± 0.061), but not GE and G_HP_E mice (*Figure 5Di*). Mirroring the weaker hippocampal drive, the light stimulation of HP augmented the power of prefrontal oscillations in theta (0.373 ± 0.094) and beta (0.269 ± 0.052) range only in CON, whereas the increase was smaller, if any, for GE (4-12Hz: 0.061 ± 0.0750, p=0.0009; 12-30Hz: 0.0640 ± 0.048, *p*=0.0009, one-way ANOVA followed by Bonferroni-corrected post hoc test) and G_HP_E (4-12Hz: 0.160 ± 0.044, p=0.007; 12-30Hz: 0.144 ± 0.040, *p*=0.003, one-way ANOVA followed by Bonferroni-corrected post hoc test) mice (*Figure 5Dii*). Similar to neonatal age, these functional deficits were related to abnormal morphology and connectivity of CA1 neurons. Detailed histological examination of the cytoarchitecture revealed that at P20 the dendritic branching of hippocampal pyramidal neurons in GE and G_HP_E mice was still significantly reduced when compared to CON (n=17~19 neurons from 3 mice in each group) mice (condition effect, *p*=7.30*10^-9^) (*Figure 5E*). These deficits were particularly prominent within a radius of 40-150 μm from the cell soma center. The sparsification of dendritic projections was accompanied by lower density of the dendritic spines in G_HP_E and GE mice when compared with CON mice (*Figure 5F*). The magnitude of density reduction was similar for basal dendrites (F(2, 52)=46.36, *p*=2.77*10^-12^, one-way ANOVA), proximal oblique dendrites (F(2, 54)=31.81, *p*=7.44*10^-10^, one-way ANOVA), and secondary apical dendrites (F(2, 53)=20.22, *p*=2.97*10^-7^, one-way ANOVA).

Taken together, these data show that the prefrontal-hippocampal dysfunction resulting from synaptic and projection deficits of CA1 pyramidal neurons in GE and G_HP_E mice persists until pre-juvenile age.

The developmental prefrontal-hippocampal dysfunction as result of hippocampal DISC1 suppression in immune-challenged mice might cause behavioral disabilities. Already at juvenile age, rodents have reliable novelty detection and recognition memory that rely on the mouse’s intrinsic exploratory drive and require no prior training or deprivation (Kruger et al., 2012). These abilities have been shown to involve communication within a circuit centered on PFC and HP (Warburton and Brown, 2015). Both GE mice and immune-challenged mice with DISC1 suppression confined to PFC have been reported to have poor recognition memory (Hartung et al., 2016; Xu et al., 2019). To identify the consequences of neonatal HP-restricted DISC1 knock-down on cognitive abilities, we tested the novel object recognition (NOR) and recency recognition (RR) in CON (n=11), G_HP_E (n=13), and GE (n=12) mice using a custom-designed arena (*Figure 6A*) and previously established protocols (*Figure 6Bi and Ci*). During the familiarization trials of these tests, all mice spent equal time investigating the two objects placed in the arena. During the NOR test trial protocols, CON mice spent significantly (*p*=2.38*10^-5^, paired t-test) longer time interacting with the novel object (79.12 ± 4.49%) than with the familiar one (20.88 ± 4.49%) (*Figure 6Bii*). In line with previous results, GE mice failed to distinguish between the two objects (familiar: 47.46 ± 7.90%; novel: 52.54 ± 7.90%, *p*=0.372, paired t-test) (*Figure 6Bii*). Similarly, pre-juvenile G_HP_E mice were also unable to distinguish between the two objects during test trial (familiar: 39.23 ± 1.29%; novel: 60.77 ± 2.19%, *p*=0.07, pared t-test) (*Figure 6Bii*). Correspondingly, the discrimination ratio between the familiar and the novel object significantly decreased in GE (0.0501 ± 0.158, *p*=0.02, one-way ANOVA followed by Bonferroni-corrected post-hoc test) and G_HP_E mice (−0.215 ± 0.142, *p*=0.02, one-way ANOVA followed by Bonferroni-corrected post-hoc test) compared with CON mice (0.582 ± 0.090) (*Figure 6Biii*).

**Figure 6.**
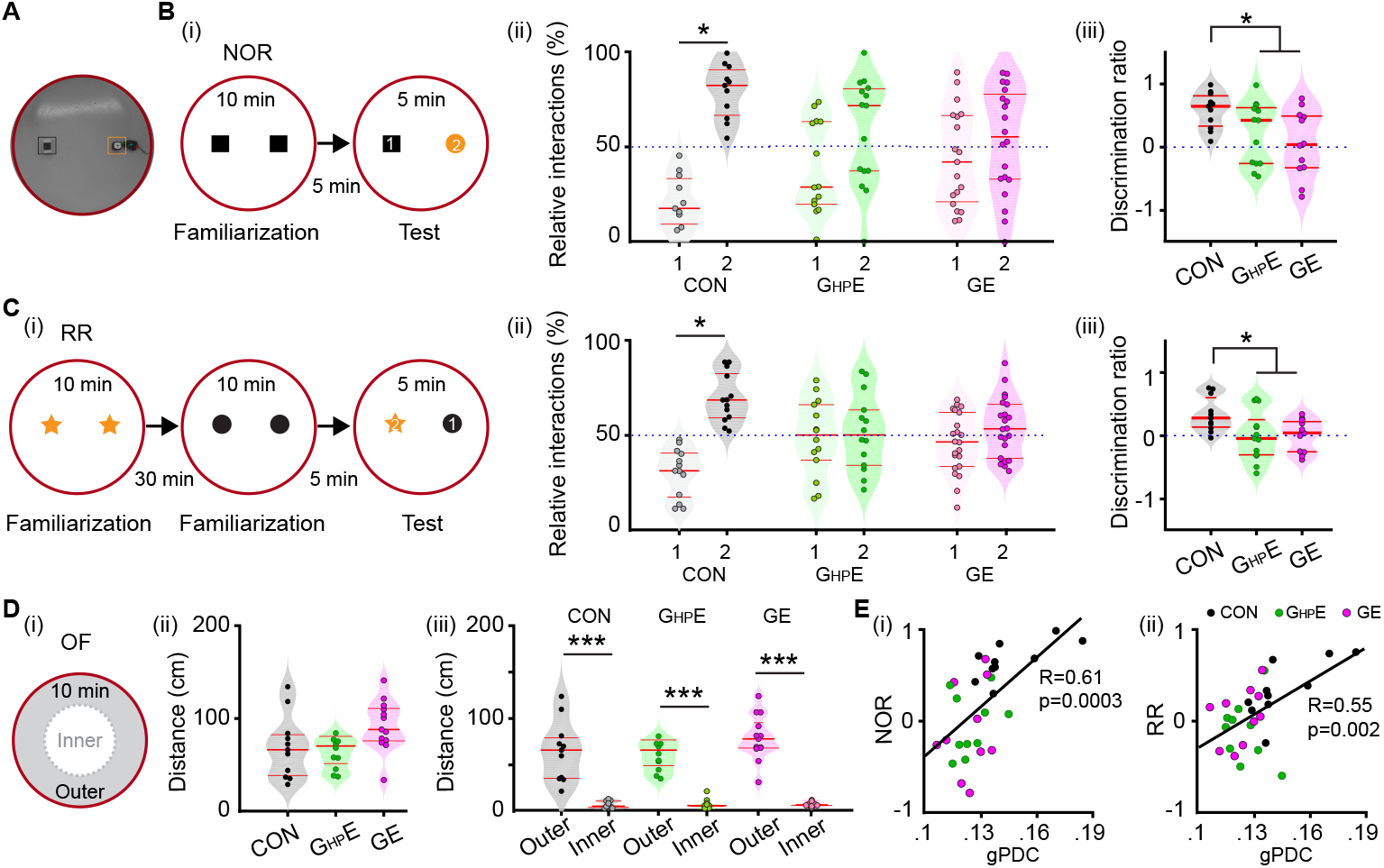
Novelty recognition of immune challenged mice with whole-brain of HP-confined DISC1 suppression. **(A)** Photograph of the arena used for NOR and RR tasks. **(B) (i)** Schematic diagrams of the protocol for NOR task. **(ii)** Violin plots displaying the relative interaction time spent by CON, G_HP_E, and GE mice with the objects during the NOR test trial. The dotted line indicates chance level. **(iii)** Violin plots displaying the NOR discrimination ratio when averaged for CON, G_HP_E and GE mice. **(C) (i)** Schematic diagrams of the protocol for RR task. **(ii)** Violin plots displaying the relative interaction time spent by CON, G_HP_E, and GE mice with the objects during the RR test trial. The dotted line indicates chance level. **(iii)** Violin plots displaying the RR discrimination ratio when averaged for CON, G_HP_E and GE mice. **(D) (i)** Schematic diagrams of the protocol for OF task. **(ii)** Violin plots displaying the distance covered in 10 mins by CON, G_HP_E, and GE mice during the OF task. **(iii)** Violin plots displaying the distance covered in the outer circle and the inner circle by CON, G_HP_E, and GE mice during OF task. **(E) (i)** The Pearson’s correlation between gPDC and discrimination ratio in NOR task. **(ii)** Same as (i) for Pearson’s correlation between gPDC and RR. Single data points are represented as dots and the red horizontal bars in violin plots represent the median and the 25^th^ and 75^th^ percentiles. *p<0.05, ***p<0.001.

During RR task, mice process temporal information by recognizing the object with which they most recently interacted. The CON mice spent more time with the object they explored during the first familiarization trial and less time with the more recent object from the second familiarization trial (old: 67.52 ± 4.72%, recent: 32.48 ± 4.72%, *p*=0.0015, paired t-test) (*Figure 6Cii*). Both G_HP_E and GE mice failed to recognize the most recently explored object and spent equal time with both objects (GE, old: 50.88 ± 3.92%, recent: 49.12 ± 3.92%, *p*=0.409, paired t-test; G_HP_E, old: 50.86 ± 5.93%, recent: 49.14 ± 5.93%, *p*=0.441, paired t-test) (*Figure 6Cii*). Correspondingly, the discrimination ratio significantly decreased in GE (0.001 ± 0.075, *p*=0.04, one-way ANOVA followed by Bonferroni-corrected post-hoc test) and G_HP_E (−0.0053 ± 0.1133, *p*=0.03, one-way ANOVA followed by Bonferroni-corrected post-hoc test) compared with the values for CON mice (0.335 ± 0.087) (*Figure 6Ciii*).

The poor performance in NOR and RR tasks may result from poor motor abilities and/or enhanced anxiety when interacting with the objects. To test this hypothesis, we analyzed the exploratory behavior of P16 CON (n=11), G_HP_E (n=13), and GE (n=12) mice in OF task (*Figure 6Di*). The distance covered in 10 mins was similar in all groups (CON: 69.176 ± 10.70 cm; GE: 77.41 ± 12.11 cm; G_HP_E: 92.25 ± 7.86 cm, F(2, 33)=1.437, *p*=0.252, one-way ANOVA) (*Figure 6Dii*). Moreover, the distance covered in the outer circle was much larger than in the inner circle of the arena (CON: 64.67 ± 10.18 cm vs. 4.50 ± 1.18 cm; GE: 72.62 ± 10.81 cm vs. 4.78 ± 1.51 cm; G_HP_E: 79.23 ± 7.00 cm vs. 5.02 ± 0.73 cm) (*Figure 6Diii*). These results indicate that exploratory and anxiety abilities were similar in CON, G_HP_E, and GE mice and thus, did not affect the recognition memory in NOR and RR tasks.

To better link the behavioral deficits in GE and G_HP_E mice with the abnormal prefrontal-hippocampal coupling, we performed the Pearson’s correlation between gPDC and discrimination ratio in NOR and RR task (*Figure 6E*). The strong positive correlation (gPDC and NOR: R= 0.61, p = 0.0003; gPDC and RR: R=0.55, p=0.002, Pearson’s correlation) confirmed that abnormal prefrontal-hippocampal coupling as results of HP-restricted DISC1 suppression in immune challenged mice might lead to poorer recognition memory at pre-juvenile age.

## Discussion

Developmental miswiring of the brain has been hypothesized to account for cognitive impairment in mental disorders. Previous studies provided first experimental evidence that the communication between PFC and HP, the core of a complex network underlying mnemonic and executive processing, is substantial impaired in mouse models of disease already at neonatal age (Hartung et al., 2016; Oberlander et al., 2019). The mechanisms causing diminished communication within prefrontal-hippocampal networks remain largely unknown. We recently identified spine loss and sparsification of dendritic projections in layer 2/3 pyramidal neurons of neonatal PFC as one mechanism of disorganized network activity and reduced coupling with HP (Chini et al., 2019; Xu et al., 2019). It is still unclear whether hippocampal dysfunction contributes to the early miswiring as well. Here, we combine electrophysiology and optogenetics *in vivo* with neuroanatomy and behavioral testing of immune challenged mice with either brain-wide or HP-confined suppression of DISC1. We provide evidence that (i) highly fragmented oscillatory activity with reduced power, fewer SPWs and decreased SPW-related firing of CA1 neurons in the i/vHP are present in both GE and G_HP_E mice; (ii) confinement of DISC1 knock-down to HP of immune challenged mice causes weaker hippocampal drive to PFC and consequently, abnormal prefrontal network activity, despite non-affected firing rates over cortical layers; (iii) HP-confined or brain-wide DISC1 suppression similarly impairs the morphology of CA1 neurons, reducing the dendritic branching and the density of spines; (iv) the morphological and functional deficits in the HP of GE and G_HP_E mice persist until pre-juvenile age leading to cognitive impairment and long-lasting disruption of underlying prefrontal-hippocampal communication.

The severe morphological and functional impairment of the developing HP when DISC1 is suppressed goes in line with previous studies that identified this gene as a hub of maturational processes (Miyoshi et al., 2003; Duan et al., 2007). Especially DISC1 knock-down in HP has been associated with long-lasting deficits and behavioral impairment related to mental disorders (Callicott et al., 2005; Meyer and Morris, 2008). DISC1 controls neurite growth, neuronal migration and differentiation as well as axon targeting (Narayan et al., 2013). In line with this function, the hippocampal structure and neuronal distribution across layers in the i/vHP was disturbed in GE mice when compared to controls. Intriguingly, HP-confined DISC1 suppression did not perturb the migration of CA1 pyramidal neurons in the dorsal HP but hinders the migration of dentate gyrus granule cells (Meyer and Morris, 2009). The arborization and synaptic interactions of CA1 pyramidal neurons seem to be profoundly altered. The sparsification of their dendritic branching and the low number of spines document major developmental deficits of CA1 neurons when DISC1 was locally suppressed. They might result from disorganized microtubule-associated dynein motor complex (Ozeki et al., 2003; Kamiya et al., 2005). Moreover, lower density of axonal projections might link HP to PFC, as previously observed for GE mice (Xu and Hanganu-Opatz, unpublished observations). The abnormal morphology is likely to underlie the early dysfunction of activity patterns generated within CA1 (e.g. SPWs, beta-gamma oscillations) and the diminishment of excitatory drive to PFC.

Suppression of DISC1 decreased the occurrence of SPWs and the SPW-related neuronal firing. While SPWs have been extensively characterized in the adult HP (Buzsaki, 1986), their underlying mechanisms during development are still largely unknown. SPWs emerge early in life (Nowack et al., 1989; Brockmann et al., 2011; Valeeva et al., 2019). Similar to adult one, the neonatal SPWs seem to be generated within HP following population bursts in CA3 area. They correlate with increased firing rate of hippocampal neurons. The decreased SPWs occurrence and related neuronal discharge indicate that DISC1 suppression might cause miswiring within HP and abnormal coupling between CA1 and CA3. Accordingly, both pulse and ramp stimuli induced hippocampal firing. However, the power of spike-triggered LFP in CA1 dramatically decreased in GE and G_HP_E mice, reflecting a weaker entrainment of local circuits in HP triggered by neuronal firing. Correspondingly, the power of oscillations in beta and gamma frequency range decreased too in GE and G_HP_E mice. In contrast, the theta bursts are less affected, their occurrence being similar across all investigated mice. This might be due to the fact that theta bursts have a multiple mostly extra-hippocampal origin with the septum as one main generator (Janiesch et al., 2011). At adulthood, the SPW-ripple events are still perturbed in different strains with suppressed DISC1, yet their occurrence was higher when compared to controls due to dysfunction of parvalbumin-positive interneurons (Altimus et al., 2015). We propose that the abnormal maturation of hippocampal circuits might have detrimental effects on the interneuron function and cause overcompensation resulting in hippocampal hyperexcitability.

Several lines of evidence show that DISC1 suppression perturbs not only the hippocampal activity and oscillatory entrainment but also the prefrontal activity and the communication within prefrontal-hippocampal networks. First, even when confined to HP, DISC1 suppression led to disorganized prefrontal activity with weaker power. In contrast, the overall firing of neurons in the PL remained unaffected. In contrast, when the DISC1 suppression was restricted to PFC, the firing of these neurons was dramatically decreased (Xu et al., 2019). Second, the timing of prelimbic firing in G_HP_E mice was disturbed by the HP-confined DISC1 suppression, the characteristic beta band peak of firing interval and HP-spike triggered LFP power in layer 2/3 of PFC being absent in these mice. Third, the synchrony over a wide frequency range and the directionality of prefrontal-hippocampal interactions diminished in GE and G_HP_E mice when compared to controls. Moreover, during ramp light stimulation the coherence of prefrontal-hippocampal interactions within beta frequency range was significantly decreased in both GE and G_HP_E mice, reflecting the weaker excitatory drive from the HP to PFC.

We propose that two mechanisms contribute to the abnormal prefrontal-hippocampal communication of GE and G_HP_E. On the one hand, due to synaptic deficits, the CA1 pyramidal neurons lose to a large extent the ability to fire in an oscillatory phase-coordinated manner. Consequently, the excitatory drive reaching mainly layer 5/6 of PFC (Parent et al., 2010; Padilla-Coreano et al., 2016) decreases and the boosting of intracortical connectivity resulting into beta entrainment within layer 2/3 is weaker. On the other hand, the sparsification of dendritic projections might cause less dense connections to the PFC. Whether the glutamate release of the hippocampal terminals targeting the PFC is also impaired remains to be elucidated.

In contrast to the DISC1 suppression confined to the pyramidal neurons in PFC, the DISC1 suppression in HP is not transient during development but persisted until pre-juvenile age. These differences across areas might result from different regulation of Disc1 gene by external cues. Over 50 proteins interact with DISC1 controlling different maturational processes (Camargo et al., 2007; Ye et al., 2017). Despite similar DISC1 knock-down by shRNA, these interactions may lead to major differences across brain regions and over time. The interference of DISC1 with immune-relevant signalling pathways is of particular relevance (Beurel et al., 2010). The structural and functional deficits caused by the combination of DISC1 suppression with MIA are by far more pronounced than the effects induced by either of the two factors. They persist throughout the life span, leading to altered social and cognitive behaviour (Abazyan et al., 2010; Ibi et al., 2010; Lipina et al., 2013). In the present study, we identified morphological and functional deficits within prefrontal-hippocampal networks of G_HP_E mice until pre-juvenile age. This abnormal communication between the two brain areas throughout the development relates to impaired pre-juvenile cognitive performance, which requires the prefrontal-hippocampal activation (Barker and Warburton, 2011). The ability to recognize new objects and their recency was absent in GE and G_HP_E mice.

While comprehensive genome-wide association studies (GWAS) showed that DISC1 is unlikely to be a “genetic” factor causing schizophrenia (Schizophrenia Working Group of the Psychiatric Genomics Consortium, 2014), a wealth of data documented its relevance for psychiatric conditions (Tomoda et al., 2016; Trossbach et al., 2016; Tomoda et al., 2017; Kakuda et al., 2019; Sawa, 2019). Orchestrating molecular cascades hypothesized to underlie disease-relevant physiological and behavioral abnormalities (Cuthbert and Insel, 2013), DISC1 points out the contribution of abnormal development for multiple mental conditions. The present results provide first insight into the mechanisms by which DISC1 suppression interferes with the circuit function and cognitive abilities. They show that besides prefrontal deficits of layer 2/3 pyramidal neurons, the dysfunction of hippocampal CA1 neurons unable to drive the down-stream PFC during early development causes impaired prefrontal-hippocampal communication that relates to poor cognitive performance. HP plays a key role for memory deficits in mental disorders (Chen et al., 2018). Impairment of hippocampal recruitment during memory tasks has been described for schizophrenia patients and high-risks subjects (Di Giorgio et al., 2013; Rasetti et al., 2014). We provide experimental evidence that developmental miswiring in hippocampus might cause cognitive deficits at adulthood and disrupt the prefrontal-hippocampal coupling. Weaker co-activation of PFC and HP has been identified in schizophrenia patients during cognitive tasks (Meyer-Lindenberg et al., 2001). Thus, the present study supports the neurodevelopmental origin of schizophrenia and highlights the hub function of hippocampus during early maturation for the functional and cognitive deficits at adulthood.

## Author contributions

I.L.H.-O., X.X. designed the experiments, X.X., L.S. carried out the experiments, X.X. analyzed the data, I.L.H.-O. and X.X. interpreted the data and wrote the paper. All authors discussed and commented on the manuscript.

## Acknowledgements

We thank Dr. Joseph Gogos for providing the DISC1 mice and Dr. A. Sawa for providing shorthairpin RNA (shRNA) to DISC1. We also thank A. Marquardt, C. Tietze, A. Dahlmann and P. Putthoff for excellent technical assistance. This work was funded by grants from the European Research Council (ERC-2015-CoG 681577 to I.L.H.-O.) and the German Research Foundation (SPP 1665, SFB 936 B5 to I.L.H.-O).

I.L. H.-O. is funding member of FENS Kavli Network of Excellence.

## References

Abazyan B, Nomura J, Kannan G, Ishizuka K, Tamashiro KL, Nucifora F, Pogorelov V, Ladenheim B, Yang C, Krasnova IN, Cadet JL, Pardo C, Mori S, Kamiya A, Vogel MW, Sawa A, Ross CA, Pletnikov MV (2010) Prenatal interaction of mutant DISC1 and immune activation produces adult psychopathology. Biol Psychiatry 68:1172–1181.

Ahlbeck J, Song L, Chini M, Bitzenhofer SH, Hanganu-Opatz IL (2018) Glutamatergic drive along the septo-temporal axis of hippocampus boosts prelimbic oscillations in the neonatal mouse. Elife 7:e33158.

Altimus C, Harrold J, Jaaro-Peled H, Sawa A, Foster DJ (2015) Disordered ripples are a common feature of genetically distinct mouse models relevant to schizophrenia. Mol Neuropsychiatry 1:52–59.

Baccala LA, Sameshima K (2001) Partial directed coherence: a new concept in neural structure determination. Biol Cybern 84:463–474.

Baccala LA, Sameshima K, Takahashi DY (2007) Generalized Partial Directed Coherence. In: 2007 15th International Conference on Digital Signal Processing, pp 163–166.

Backus AR, Schoffelen JM, Szebenyi S, Hanslmayr S, Doeller CF (2016) Hippocampal-Prefrontal Theta Oscillations Support Memory Integration. Curr Biol 26:450–457.

Barker GR, Warburton EC (2011) When is the hippocampus involved in recognition memory? J Neurosci 31:10721–10731.

Baumgart J, Grebe N (2015) C57BL/6-specific conditions for efficient in utero electroporation of the central nervous system. J Neurosci Methods 240:116–124.

Berndt A, Schoenenberger P, Mattis J, Tye KM, Deisseroth K, Hegemann P, Oertner TG (2011) High-efficiency channelrhodopsins for fast neuronal stimulation at low light levels. PNAS 108:7595–7600.

Beurel E, Michalek SM, Jope RS (2010) Innate and adaptive immune responses regulated by glycogen synthase kinase-3 (GSK3). Trends Immunol 31:24–31.

Bitzenhofer SH, Hanganu-Opatz IL (2014) Oscillatory coupling within neonatal prefrontal-hippocampal networks is independent of selective removal of GABAergic neurons in the hippocampus. Neuropharmacology 77:57–67.

Bitzenhofer SH, Ahlbeck J, Hanganu-Opatz IL (2017a) Methodological Approach for Optogenetic Manipulation of Neonatal Neuronal Networks. Front Cell Neurosci 11:239.

Bitzenhofer SH, Ahlbeck J, Wolff A, Wiegert JS, Gee CE, Oertner TG, Hanganu-Opatz IL (2017b) Layer-specific optogenetic activation of pyramidal neurons causes beta-gamma entrainment of neonatal networks. Nat Commun 8:14563.

Brockmann MD, Poschel B, Cichon N, Hanganu-Opatz IL (2011) Coupled oscillations mediate directed interactions between prefrontal cortex and hippocampus of the neonatal rat. Neuron 71:332–347.

Buzsaki G (1986) Hippocampal Sharp Waves - Their Origin and Significance. Brain Research 398:242–252.

Buzsaki G (2002) Theta oscillations in the hippocampus. Neuron 33:325–340.

Callicott JH, Straub RE, Pezawas L, Egan MF, Mattay VS, Hariri AR, Verchinski BA, Meyer-Lindenberg A, Balkissoon R, Kolachana B, Goldberg TE, Weinberger DR (2005) Variation in DISC1 affects hippocampal structure and function and increases risk for schizophrenia. PNAS 102:8627–8632.

Camargo LM, Collura V, Rain JC, Mizuguchi K, Hermjakob H, Kerrien S, Bonnert TP, Whiting PJ, Brandon NJ (2007) Disrupted in Schizophrenia 1 Interactome: evidence for the close connectivity of risk genes and a potential synaptic basis for schizophrenia. Mol Psychiatry 12:74–86.

Cash-Padgett T, Jaaro-Peled H (2013) DISC1 mouse models as a tool to decipher gene-environment interactions in psychiatric disorders. Front Behav Neurosci 7:113.

Chen LW, Sun D, Davis SL, Haswell CC, Dennis EL, Swanson CA, Whelan CD, Gutman B, Jahanshad N, Iglesias JE, Thompson P, Mid-Atlantic MW, Wagner HR, Saemann P, LaBar KS, Morey RA (2018) Smaller hippocampal CA1 subfield volume in posttraumatic stress disorder. Depress Anxiety 35:1018–1029.

Chini M, Popplau JA, Lindemann C, Carol-Perdiguer L, Hnida M, Oberlander V, Xu X, Ahlbeck J, Bitzenhofer SH, Mulert C, Hanganu-Opatz IL (2019) Resolving and Rescuing Developmental Miswiring in a Mouse Model of Cognitive Impairment. Neuron.

Cichon NB, Denker M, Grun S, Hanganu-Opatz IL (2014) Unsupervised classification of neocortical activity patterns in neonatal and pre-juvenile rodents. Front Neural Circuits 8:50.

Clapcote SJ, Lipina TV, Millar JK, Mackie S, Christie S, Ogawa F, Lerch JP, Trimble K, Uchiyama M, Sakuraba Y, Kaneda H, Shiroishi T, Houslay MD, Henkelman RM, Sled JG, Gondo Y, Porteous DJ, Roder JC (2007) Behavioral phenotypes of Disc1 missense mutations in mice. Neuron 54:387–402.

Clement EA, Richard A, Thwaites M, Ailon J, Peters S, Dickson CT (2008) Cyclic and sleep-like spontaneous alternations of brain state under urethane anaesthesia. PLoS One 3:e2004.

Crabtree GW, Sun Z, Kvajo M, Broek JA, Fenelon K, McKellar H, Xiao L, Xu B, Bahn S, O’Donnell JM, Gogos JA (2017) Alteration of Neuronal Excitability and Short-Term Synaptic Plasticity in the Prefrontal Cortex of a Mouse Model of Mental Illness. J Neurosci 37:4158–4180.

Cuthbert BN, Insel TR (2013) Toward the future of psychiatric diagnosis: the seven pillars of RDoC. BMC Med 11:126.

Di Giorgio A, Gelao B, Caforio G, Romano R, Andriola I, D’Ambrosio E, Papazacharias A, Elifani F, Bianco LL, Taurisano P, Fazio L, Popolizio T, Blasi G, Bertolino A (2013) Evidence that hippocampal-parahippocampal dysfunction is related to genetic risk for schizophrenia. Psychol Med 43:1661–1671.

Duan X, Chang JH, Ge S, Faulkner RL, Kim JY, Kitabatake Y, Liu XB, Yang CH, Jordan JD, Ma DK, Liu CY, Ganesan S, Cheng HJ, Ming GL, Lu B, Song H (2007) Disrupted-In-Schizophrenia 1 regulates integration of newly generated neurons in the adult brain. Cell 130:1146–1158.

Eichenbaum H (2017) Prefrontal-hippocampal interactions in episodic memory. Nat Rev Neurosci 18:547–558.

Ennaceur A, Delacour J (1988) A new one-trial test for neurobiological studies of memory in rats. 1: Behavioral data. Behav Brain Res 31:47–59.

Hartung H, Cichon N, De Feo V, Riemann S, Schildt S, Lindemann C, Mulert C, Gogos JA, Hanganu-Opatz IL (2016) From Shortage to Surge: A Developmental Switch in Hippocampal-Prefrontal Coupling in a Gene-Environment Model of Neuropsychiatric Disorders. Cereb Cortex 26:4265–4281.

Heyser CJ, Ferris JS (2013) Object exploration in the developing rat: methodological considerations. Dev Psychobiol 55:373–381.

Ibi D, Nagai T, Koike H, Kitahara Y, Mizoguchi H, Niwa M, Jaaro-Peled H, Nitta A, Yoneda Y, Nabeshima T, Sawa A, Yamada K (2010) Combined effect of neonatal immune activation and mutant DISC1 on phenotypic changes in adulthood. Behav Brain Res 206:32–37.

Janiesch PC, Kruger HS, Poschel B, Hanganu-Opatz IL (2011) Cholinergic control in developing prefrontal-hippocampal networks. J Neurosci 31:17955–17970.

Kakuda K, Niwa A, Honda R, Yamaguchi KI, Tomita H, Nojebuzzaman M, Hara A, Goto Y, Osawa M, Kuwata K (2019) A DISC1 point mutation promotes oligomerization and impairs information processing in a mouse model of schizophrenia. J Biochem 165:369–378.

Kamiya A, Kubo K, Tomoda T, Takaki M, Youn R, Ozeki Y, Sawamura N, Park U, Kudo C, Okawa M, Ross CA, Hatten ME, Nakajima K, Sawa A (2005) A schizophrenia-associated mutation of DISC1 perturbs cerebral cortex development. Nat Cell Biol 7:1167–1178.

Koike H, Arguello PA, Kvajo M, Karayiorgou M, Gogos JA (2006) Disc1 is mutated in the 129S6/SvEv strain and modulates working memory in mice. PNAS 103:3693–3697.

Kruger HS, Brockmann MD, Salamon J, Ittrich H, Hanganu-Opatz IL (2012) Neonatal hippocampal lesion alters the functional maturation of the prefrontal cortex and the early cognitive development in pre-juvenile rats. Neurobiol Learn Mem 97:470–481.

Kvajo M, McKellar H, Arguello PA, Drew LJ, Moore H, MacDermott AB, Karayiorgou M, Gogos JA (2008) A mutation in mouse Disc1 that models a schizophrenia risk allele leads to specific alterations in neuronal architecture and cognition. PNAS 105:7076–7081.

Kvajo M, McKellar H, Drew LJ, Lepagnol-Bestel AM, Xiao L, Levy RJ, Blazeski R, Arguello PA, Lacefield CO, Mason CA, Simonneau M, O’Donnell JM, MacDermott AB, Karayiorgou M, Gogos JA (2011) Altered axonal targeting and short-term plasticity in the hippocampus of Disc1 mutant mice. PNAS 108:1349–1358.

Lipina TV, Zai C, Hlousek D, Roder JC, Wong AH (2013) Maternal immune activation during gestation interacts with Disc1 point mutation to exacerbate schizophrenia-related behaviors in mice. J Neurosci 33:7654–7666.

Meyer-Lindenberg A, Poline JB, Kohn PD, Holt JL, Egan MF, Weinberger DR, Berman KF (2001) Evidence for abnormal cortical functional connectivity during working memory in schizophrenia. Am J Psychiatry 158:1809–1817.

Meyer KD, Morris JA (2008) Immunohistochemical analysis of Disc1 expression in the developing and adult hippocampus. Gene Expr Patterns 8:494–501.

Meyer KD, Morris JA (2009) Disc1 regulates granule cell migration in the developing hippocampus. Hum Mol Genet 18:3286–3297.

Miyoshi K, Honda A, Baba K, Taniguchi M, Oono K, Fujita T, Kuroda S, Katayama T, Tohyama M (2003) Disrupted-In-Schizophrenia 1, a candidate gene for schizophrenia, participates in neurite outgrowth. Mol Psychiatry 8:685–694.

Narayan S, Nakajima K, Sawa A (2013) DISC1: a key lead in studying cortical development and associated brain disorders. Neuroscientist 19:451–464.

Nowack WJ, Janati A, Angtuaco T (1989) Positive temporal sharp waves in neonatal EEG. Clin Electroencephalogr 20:196–201.

Oberlander VC, Xu X, Chini M, Hanganu-Opatz IL (2019) Developmental dysfunction of prefrontal-hippocampal networks in mouse models of mental illness. Eur J Neurosci 50:3072–3084.

Ozeki Y, Tomoda T, Kleiderlein J, Kamiya A, Bord L, Fujii K, Okawa M, Yamada N, Hatten ME, Snyder SH, Ross CA, Sawa A (2003) Disrupted-in-Schizophrenia-1 (DIsC-1): mutant truncation prevents binding to NudE-like (NUDEL) and inhibits neurite outgrowth. PNAS 100:289–294.

Padilla-Coreano N, Bolkan SS, Pierce GM, Blackman DR, Hardin WD, Garcia-Garcia AL, Spellman TJ, Gordon JA (2016) Direct Ventral Hippocampal-Prefrontal Input Is Required for Anxiety-Related Neural Activity and Behavior. Neuron 89:857–866.

Parent MA, Wang L, Su J, Netoff T, Yuan LL (2010) Identification of the hippocampal input to medial prefrontal cortex in vitro. Cereb Cortex 20:393–403.

Parks DH, Rinke C, Chuvochina M, Chaumeil PA, Woodcroft BJ, Evans PN, Hugenholtz P, Tyson GW (2017) Recovery of nearly 8,000 metagenome-assembled genomes substantially expands the tree of life. Nat Microbiol 2:1533–1542.

Rasetti R, Mattay VS, White MG, Sambataro F, Podell JE, Zoltick B, Chen Q, Berman KF, Callicott JH, Weinberger DR (2014) Altered hippocampal-parahippocampal function during stimulus encoding: a potential indicator of genetic liability for schizophrenia. JAMA Psychiatry 71:236–247.

Ripke S et al. (2013) Genome-wide association analysis identifies 13 new risk loci for schizophrenia. Nat Genet 45:1150–1159.

Sawa A (2019) DISC1 and Its Protein Interactomes for Mental Function. Biol Psychiatry 85:283–284.

Siapas AG, Lubenov EV, Wilson MA (2005) Prefrontal phase locking to hippocampal theta oscillations. Neuron 46:141–151.

Spellman T, Rigotti M, Ahmari SE, Fusi S, Gogos JA, Gordon JA (2015) Hippocampal-prefrontal input supports spatial encoding in working memory. Nature 522:309–314.

Stark KL, Xu B, Bagchi A, Lai WS, Liu H, Hsu R, Wan X, Pavlidis P, Mills AA, Karayiorgou M, Gogos JA (2008) Altered brain microRNA biogenesis contributes to phenotypic deficits in a 22q11-deletion mouse model. Nat Genet 40:751–760.

Stujenske JM, Spellman T, Gordon JA (2015) Modeling the Spatiotemporal Dynamics of Light and Heat Propagation for In Vivo Optogenetics. Cell Rep 12:525–534.

Sullivan PF, Daly MJ, O’Donovan M (2012) Genetic architectures of psychiatric disorders: the emerging picture and its implications. Nat Rev Genet 13:537–551.

Szczurkowska J, Cwetsch AW, dal Maschio M, Ghezzi D, Ratto GM, Cancedda L (2016) Targeted in vivo genetic manipulation of the mouse or rat brain by in utero electroporation with a triple-electrode probe. Nat Protoc 11:399–412.

Tomoda T, Hikida T, Sakurai T (2017) Role of DISC1 in Neuronal Trafficking and its Implication in Neuropsychiatric Manifestation and Neurotherapeutics. Neurotherapeutics 14:623–629.

Tomoda T, Sumitomo A, Jaaro-Peled H, Sawa A (2016) Utility and validity of DISC1 mouse models in biological psychiatry. Neuroscience 321:99–107.

Trossbach SV et al. (2016) Misassembly of full-length Disrupted-in-Schizophrenia 1 protein is linked to altered dopamine homeostasis and behavioral deficits. Mol Psychiatry 21:1561–1572.

Valeeva G, Nasretdinov A, Rychkova V, Khazipov R (2019) Bilateral Synchronization of Hippocampal Early Sharp Waves in Neonatal Rats. Front Cell Neurosci 13:29.

Warburton EC, Brown MW (2015) Neural circuitry for rat recognition memory. Behav Brain Res 285:131–139.

Wolansky T, Clement EA, Peters SR, Palczak MA, Dickson CT (2006) Hippocampal slow oscillation: a novel EEG state and its coordination with ongoing neocortical activity. J Neurosci 26:6213–6229.

Xu X, Chini M, Bitzenhofer SH, Hanganu-Opatz IL (2019) Transient Knock-Down of Prefrontal DISC1 in Immune-Challenged Mice Causes Abnormal Long-Range Coupling and Cognitive Dysfunction throughout Development. J Neurosci 39:1222–1235.

Ye F, Kang E, Yu C, Qian X, Jacob F, Yu C, Mao M, Poon RYC, Kim J, Song H, Ming GL, Zhang M (2017) DISC1 Regulates Neurogenesis via Modulating Kinetochore Attachment of Ndel1/Nde1 during Mitosis. Neuron 96:1041–1054e1045.

